# Dual-Engineered Dendritic Cell–Derived Small Extracellular Vesicles Couple T-Cell Priming with Checkpoint Reprogramming for Synergistic Immunotherapy

**DOI:** 10.64898/2026.04.08.717283

**Authors:** Gaeun Kim, Shiyu Wang, Runyao Zhu, Matthew J. Webber, Xin Lu, Yichun Wang

## Abstract

Immunotherapy has transformed cancer treatment, yet cell-based therapies remain complex and costly, and immune checkpoint blockade (ICB) agents often suffer from limited stability and poor T-cell selectivity. Here, we develop an engineered dendritic cell–derived small extracellular vesicle (DC-sEV) nanoplatform for combinatorial immunotherapy via *in situ* T-cell activation and checkpoint reprogramming. DC-sEVs preserve intrinsic dendritic-cell immunobiology, enabling antigen presentation and potent T-cell activation. We further integrate high-efficiency cargo loading and membrane functionalization to selectively deliver ICB payloads to T cells, achieving dual reprogramming that sustains effector function and amplifies antitumor immunity. This approach reduced cancer cell viability to 44.05% *in vitro* and produced 82.12% tumor growth inhibition *in vivo*, establishing DC-sEVs as a targeted, scalable cell-free immunotherapy platform.

**HIGHLIGHTS:**

- DC-sEVs preserve antigen presentation, T-cell activation, and lymph node targeting
- Chirality-assisted loading with pH-responsive functionalization enables efficient cytosolic delivery while maintaining membrane bioactivity
- Engineered DC-sEVs combine in situ T-cell priming and PD-1 silencing to enhance effector function
- In situ T-cell reprogramming drives potent antitumor efficacy and favorable tumor microenvironment remodeling

**GRAPHICAL ABSTRACT:** 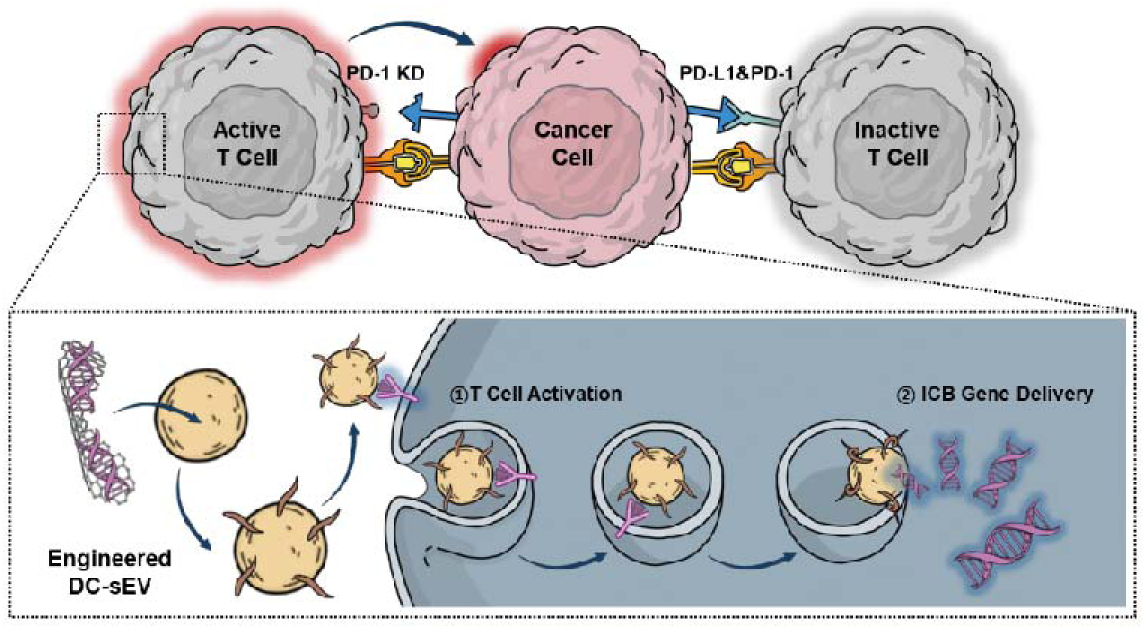

## 1. INTRODUCTION

Immunotherapy has advanced cancer treatment by mobilizing the immune system to recognize and eliminate malignant cells. Modalities such as adoptive T□cell transfer, immune checkpoint blockade (ICB), and dendritic cell (DC) vaccines have demonstrated clinical benefit across diverse malignancies.[1] Nevertheless, each approach faces challenges that restrict durable and broadly effective responses. Engineered T□cell therapies, including CAR-T cells, are labor-intensive and costly to manufacture and often show limited persistence after infusion.[1] Meanwhile, tumors suppress immune surveillance through inhibitory checkpoints such as PD-L1/PD-1, driving T-cell dysfunction and exhaustion.[2] Although ICB can reinvigorate antitumor T□cell activity,[3] its efficacy is frequently constrained by inadequate tumor delivery and suboptimal stability/targetability *in vivo*, motivating improved delivery strategies.[4] Combination regimens pairing adoptive T□cell therapy with ICB have achieved partial success, yet fully realizing their synergistic potential remains challenging.[5,6] Hence, there is a critical need for multifunctional strategies that boost T-cell function while preventing exhaustion to achieve durable antitumor immunity.

Small extracellular vesicles (sEVs) are nanoscale, lipid bilayer–enclosed vesicles (∼50–150 nm in diameter) that are secreted by most eukaryotic cells.[7] sEVs carry parent cell–derived membrane components and diverse bioactive cargos (proteins, lipids, and nucleic acids) that mediate intercellular communication.[8,9] Due to their biocompatibility and cargo-protective properties, sEVs have emerged as attractive therapeutic delivery vehicles.[10,11] In this context, immune cell–derived sEVs are of particular interest to tumor immunotherapy because they can reflect the immunological functions of their cells of origin.[12] Notably, DC–derived sEVs (DC-sEVs) have been explored as a cell-free alternative to DC vaccines,[13] as they retain key features of the parental DC membrane and can modulate immune responses by presenting antigens via inherited major histocompatibility complex (MHC) molecules together with co-stimulatory ligands to initiate adaptive immunity through interactions with T cells.[14,15] Engagement of cognate T-cell receptors (TCRs) by peptide–MHC complexes on DC-sEVs can promote vesicle–T cell interactions and facilitate uptake, thereby supporting downstream signaling events that contribute to T-cell activation.[16–18]

In this study, we leveraged the antigen-presenting capacity, T cell–interactive properties, and drug-encapsulation potential of DC-sEVs to develop a combinatorial immunotherapy (**Figure 1**). We generated MHC- and co-stimulatory ligand–enriched vesicles from mature DCs[19–21] and equipped them with PD-1–targeting small interfering RNA (siRNA; siPD-1) using a high-efficiency chirality–assisted loading strategy, while preserving membrane bioactivity. To enable productive cytosolic delivery, we further introduced the pH-responsive fusogenic peptide to promote endosomal escape and intracellular release. The resulting engineered DC-sEVs preferentially entered T cells, boosted antitumor activation programs, and delivered functional PD-1 knockdown, translating into potent tumor control with effective immune remodeling both *in vitro* and *in vivo*. Together, our work establishes engineered DC-sEVs as a versatile nanoplatform for integrating immune stimulation and gene-based checkpoint blockade to advance next-generation immunotherapy.

**Figure 1.**
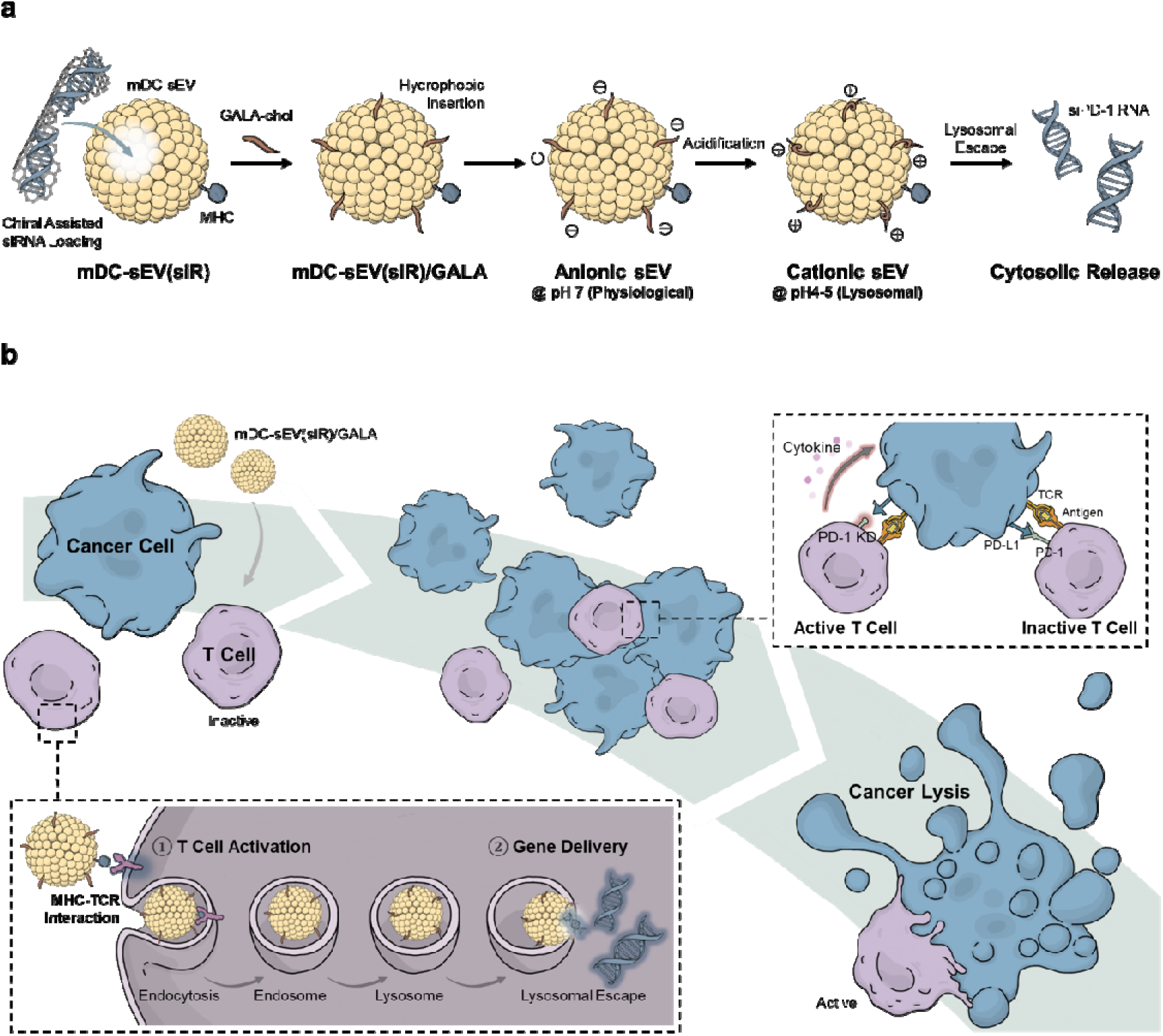
Schematic illustration of engineered mature dendritic cells (mDCs) derived small extracellular vesicles (sEVs) for *in situ* T-cell reprogramming in cancer therapy. (**a**) Engineering strategy of mDC-sEVs and (**b**) Efficient antigen presentation (MHC-mediated)–driven T-cell activation and delivery of small interfering RNA (siRNA) against the immune checkpoint receptor (PD-1) in activated T cells, resulting in boosted antitumor immune responses.

## 2. METHODS AND METHODS

### 2.1. Materials

Carbon nanofibers (719803), Hydrochloric acid (320331) and DAPI (D9542) were purchased from Sigma-Aldrich (MO, USA). Sulfuric acid (BDH3068-500MLP), Nitric acid (BDH3044-500MLPC), syringe filter (0.22 μm; 76479-010), and Fetal bovine serum (FBS; 1300-500) were purchased from VWR (PA, USA). Dialysis membrane tubing (MWCO: 1 kD; 20060186) was purchased from Spectrum Chemical Manufacturing Company (NJ, USA). 1-ethyl-3-(3-dimethyl-aminopropyl) carbodiimide (EDC; 22980), VybrantTM Multicolor Cell-Labeling Kit (DiD; V22889), Pierce BCA Protein Assay Kits (23225), GibcoTM Trypsin-EDTA (25200072), Ovalbumin (OVA; PI77120), Mouse GM-CSF Recombinant Protein (315-03-20UG, PeproTech), CellTrace CFSE Cell Proliferation Kit (C34570), OVA257-264 (SIINFEKL) peptide bound to H-2Kb Monoclonal Antibody (eBio25-D1.16 (25-D1.16), PE, eBioscience; 12-5743-82), Silencer Select Pre-Designed Mouse PD-1 siRNA (s71358) and FITC CD8a Monoclonal Antibody (53-6.7; 11-0081-82) were purchased from Thermofisher Scientific (MA, USA). N-hydroxysuccinimide sodium salt (Sulfo-NHS; 56485), and *D*-cysteine (A110205-011) were purchased from AmBeed (IL, USA). Sodium Hydroxide (1310-73-2) was purchased from Ward’s Science (NY, USANitrocellulose membrane (1662807), UView 6x Loading Dye (1665111), and Clarity Max Western Enhanced Chemiluminescence (ECL) Substrate (1705060) were purchased from Bio-Rad (CA, USA). RIPA Buffer (9806), anti-mouse Horseradish Peroxidase (HRP)-linked secondary antibody (7076), anti-rat HRP-linked secondary antibody (7077), anti-rabbit HRP-linked secondary antibody (7074), and anti-PD-1 antibody (84651S) were purchased from Cell Signaling Technology (MA, USA). Anti-beta-actin antibody (sc-47778), Anti-CD9 antibody (sc-13118), anti-CD63 antibody (sc-5275), anti-CD81 antibody (sc-166029), anti-MHC class I antibody (sc-59199), anti-MHC class II antibody (sc-32247), anti-CD80 antibody (sc-376012), anti-CD86 antibody (sc-28347), anti-CD40 antibody (sc-65263), and CD274/PD-L1 siRNA (sc-39700) were purchased from Santa Cruz Biotechnology (TX, USA). Cell Counting Kit-8 (ALX-850-039-KI01) was purchased from Enzo Biochem (NY, USA). CytoTrace Red CMTPX (22015) were purchased from AAT Bioquest (CA, USA). Cy3-labeled PD-L1 siRNA(19 bp) was purchased from BOC Sciences (NY, USA). Phosphate-buffered saline (PBS; 21-040-CM) and Minimum essential medium eagle (MEM; 10-010-CV) were purchased from Corning (NY, USA). Antibiotic antimycotic (15240096) was purchased from Fisher Scientific (MA, USA). 4% Paraformaldehyde (PFA; 15735-50S), and UranyLess (22409) were purchased from Electron Microscopy Sciences (PA, USA). Tween-20 (BTNM-0080) was purchased from G-Biosciences (MO, USA). Cryopres dimethyl sulfoxide (DMSO; 092780148) was purchased from MP Biomedicals (CA, USA). Zombie Aqua Fixable Viability Kit (423101), APC anti-mouse CD69 Antibody (104514), FITC anti-mouse CD279 (PD-1) Antibody (135214), FITC anti-mouse TNF-α Antibody (506304), PE/Cyanine7 anti-human/mouse Granzyme B Recombinant Antibody (372214), PE anti-mouse IL-2 Antibody (503808), Ultra-LEAF™ Purified anti-mouse CD28 Antibody (102116), Ultra-LEAF™ Purified anti-mouse CD3ε Antibody (100340), and MojoSort™ Mouse CD8 T Cell Isolation Kit (480008) were purchased from BioLegend (CA, USA). PE Anti-Mouse CD279 (PD-1) (J43.1) (50-9985-U025), PE Anti-Mouse IFN gamma (XMG1.2) (50-7311-U025) were purchased from Cytek Biosciences (CA, USA).

### 2.2. Instruments

Transmission electron microscopy (TEM; Talos F200i (S); Thermofisher Scientific, MA, USA) was used to confirm the nanostructures. Nanoparticle tracking analysis (NTA; NanoSight LM10 system; Malvern Instrument Ltd., Worcestershire, UK) was used to measure the size distribution and concentration of nanoparticles. Single-molecule localization microscopy (SMLM; ONI, CA, USA) was used for super-resolution microscopic images of individual sEVs. Fluorescence was measured by a plate reader (Infinite 200 PRO; Tecan, Männedorf, Switzerland). Circular dichroism (CD) spectrometer (Jasco J-1700 Spectrometer; Jasco International Company, MD, USA) was used to measure the absorbance of polarized light were measured. Bruker Tensor 27 Fourier-transform infrared (FTIR) Spectrometer (Bruker Optics International Company, MA, USA) was used to analyze the Attenuated Total Reflectance (ATR)-FTIR. Mass spectrometer (MS; Thermo Q-Exactive HF, Thermo Fisher, MA, USA) was used to identify the proteins. Western blot and agarose gel images acquired using two bioimaging systems (Azure 600; Azure Biosystems, CA, USA). Humidified incubator (MCO-15AC; Osaka, Japan) was used to maintain and incubate all cell lines used in this study. Fluorescence-activated cell sorting system (FACS; Cytek Northern LightsTM (NL)-CLC NL-2000; Cytek Biosciences, CA, USA) was used to sort and detect the cells. Flow Cytometer (FC; CytoFLEX S; Beckman Coulter, CA, USA) was used to sort and detect the cells.FACS and FC data was analyzed using FlowJo software v.10.10.0 (FlowJo LLC, OR, USA). Advanced Molecular Imaging HT Instrument (SPECTRAL AMI HT; Spectral Instruments Imaging, AZ, USA) was used for *ex vivo* mice study. Brightfield microscope (Keyence BZ-X810; Keyence Corporation, IL, USA) was used for H&E histological imaging. SigmaPlot 10.0 (Systat Software Inc., CA, USA) was used to generate all plots of the analyzed data.

### 2.3. Dendritic Cell Maturation and sEV Isolation

Immature dendritic cells, isolated from the bone marrow of a mouse (JAWS II; CRL-2612, ATCC, VA, USA), were cultured in MEM supplemented with 20% FBS, 1% antibiotic-antimycotic, and 5ng/mL GM-CSF in a humidified incubator at 37°C with 5% CO2. For the comparison, mouse fibroblast (3T3; CRL-1658, ATCC, VA, USA), were cultured in MEM supplemented with 10% FBS and 1% antibiotic-antimyotic in a humidified incubator at 37°C with 5% CO2. The cells were washed with 1× PBS, trypsinized with Trypsin-EDTA for passages. DC Maturation was induced as follows: OVA dose dependent test for optimization sEV isolation conducted as follows: Once the cells were vibrant after passage from the same batch, the culture medium was replaced with medium containing varying concentrations of the compound (0–140 µg/mL), followed by 24 hours of incubation. Cell morphology and confluency were imaged using a VWR Basic Inverted Microscope (76317-470, VWR; PA, USA) equipped with an OMAX 5.0 MP High Sensitivity CCD Digital Camera (A3550UPA-R75, OMAX; WA, USA). Western blot analysis (see Methods for details) was performed to confirm MHC expression levels in an OVA-dependent manner. The sEVs were isolated as follows: Once cells reached 70–80% confluency in the flasks, the culture media were replaced with serum-free media, followed by three washes with 1× PBS. After 24 hours of incubation, the serum-free media from all three cell types were collected and processed using vacuum filtration systems (0.22 μm pore size; 10040-460; VWR, PA, USA) to remove undesired large debris. The filtered media was then subjected to ultrafiltration using centrifugal devices (100 kDa cutoff; Spin-X UF Concentrator; Corning, NY, USA) and washed with 1× PBS buffer until transparent sEV suspension was collected. sEV characterization was performed as follows: TEM (80 kV)was imaged with negative staining by UranylLess and analyzed using ImageJ software. NTA measurements were performed using diluted sEVs (100-fold dilution; 10 μL in 1 mL of PBS buffer) and analyzed with the NTA 3.3 analytical software suite. Western blot analysis (see Methods for details) was performed to verify the expression of sEV markers (CD9, CD63, and CD81), MHC (I and II), and costimulatory ligands (CD40, CD80, CD86). OVA-specific MHC class I expression was identified on OVA-stimulated cells and their sEVs by flow cytometry using the 25-D1.16 monoclonal antibody, which specifically recognizes the ovalbumin-derived peptide SIINFEKL presented by H-2Kb.

### 2.4. sEV Engineering: Chiral graphene quantum dot (GQD)-Assisted siRNA Loading and GALA Surface Functionalization

Chiral graphene quantum dots functionalized with *D*-cysteine (*D*-GQDs) were synthesized according to previously reported methods:[22,23] *D*-GQDs were complexed with siRNA at an optimized binding ratio (*D*-GQD:siRNA = 1:2) and incubated until binding saturation was achieved (∼10 min). To load sEVs with siRNA, purified sEVs were then co-incubated with the siRNA/*D*-GQD complexes under gentle mixing at the optimized concentration reported previously (sEVs:*D*-GQDs =1 × 10^9^ particles/mL: 15 μM)[22] and incubated at room temperature until loading saturation (∼15 min). The siRNA-loaded sEVs were subsequently washed once with cold PBS (4 °C) using a 100 kDa centrifugal filter to remove loosely associated, unincorporated siRNA/*D*-GQD complexes. The pH-responsive peptide GALA conjugated with cholesterol (GALA-chol) was synthesized according to a previously reported method.[24] GALA-chol was co-incubated with siRNA-loaded sEVs at the optimized concentration (sEVs:GALA-chol = 1 × 10^9^ particles:6 μM) at room temperature for at least 10 min. Subsequently, the siRNA-loaded sEVs with GALA-functionalized (sEV(siR)/GALA) were washed four times with cold PBS (4 °C) using a 100 kDa centrifugal filter to remove unincorporated GALA-chol as well as any residual free siRNA/*D*-GQD complexes remaining from prior washing steps. The resulting engineered sEV(siR)/GALA were immediately stored at 4 °C and used within 2 h for *in vitro* and *in vivo* studies. The surface charge of sEVs was measured using a Zetasizer with the Hückel model, as optimized in our previous study.[9,24] Briefly, a ζ-potential cuvette (DTS1070, Malvern Instruments Ltd., Worcestershire, U.K.) was rinsed once with 100% ethanol and washed twice with distilled water. sEVs diluted to (2–3) × 10□ particles/mL in solutions of varying pH, adjusted with HCl or NaOH, were loaded into the cuvette. Five measurements per sample were performed with 30–100 autoruns each. All constituents used for sEV engineering, including the preparation of *D*-GQDs and GALA-chol, as well as additional characterization and optimization procedures (e.g., complexation, membrane permeability, and loading efficiency) for verification of successful engineering, are described in detail in the Supporting Information.

### 2.5. CD8□ T cell isolation for *in vitro* study

CD8□ T cells were isolated from the spleens of OT-1 mice using the Mouse CD8□ T Cell Isolation Kit (MojoSort; 480008; BioLegend, CA, USA) according to the manufacturer’s instructions. Mouse T cell culture medium for activation: Roswell Park Memorial Institute 1640 (RPMI 1640, SH30027.FS; Cytiva) + 10% FBS (SH30396.03; Cytiva) + 1% Penicillin-Streptomycin (P/S, SV30010; Cytiva) + 50 µM of 2-mercaptoethanol (Sigma-Aldrich, M3148-100ML) + 50 U/mL murine Interleukin-2 (IL-2, 575404; BioLegend).

### 2.6. Flow Cytometry Analysis

T cell proliferation was assessed as follows: First, purified CD8□ T cells were labeled with carboxyfluorescein succinimidyl ester (CFSE; Invitrogen V12883) according to the manufacturer’s protocol and co-incubated with cells or sEVs. After 4 days of incubation, cells were harvested, stained with DAPI, and washed. They were then analyzed by flow cytometry through the appropriate analysis workflow (**Figure S36**).

GALA concentration-dependent, time-dependent, and temperature-dependent sEV uptake into T cells was assessed as follows: Purified CD8□ T cells were co-incubated with *D*-GQD–loaded sEVs. For optimization of GALA functionalization, GALA-chol concentration was varied from 0–10 µM per 10^9^ sEVs and incorporated to the sEVs in accordance with our previous study.[24] With the corresponding control conditions (time and temperature), T cells were washed and then analyzed by flow cytometry through the appropriate analysis workflow (**Figure S37**).

T cell activation was assessed as follows: Purified CD8□ T cells were co-incubated with sEVs for 4 days. Cells were collected, stained with DAPI, and subsequently stained with fluorochrome-conjugated antibodies specific for surface markers such as CD69 and PD-1, followed by washing. They were then analyzed by flow cytometry through the appropriate analysis workflow (**Figure S38**).

T cell cytokine production was assessed as follows: Activated CD8□ T cells, confirmed from the T cell activation test above, underwent intracellular cytokine staining following stimulation with PMA/ionomycin (Cell Activation Cocktail, 423301; BioLegend) in the presence of brefeldin A (420601; Biolegend). After 4 h, T cells were collected, fixed and permeabilized, and stained with Zombie Aqua™ Fixable Viability Kit (423101; Biolegend) together with fluorochrome-conjugated antibodies specific for CD8, Granzyme B, IFN-γ, IL-2, and TNF-α, followed by washing. Cells were then analyzed by flow cytometry through the appropriate analysis workflow (**Figure S39**).

PD-1 silencing after siRNA delivery was assessed as follows: Purified CD8□ T cells were co-incubated with sEVs for 2 days. Cells were collected, stained with DAPI, and subsequently stained with fluorochrome-conjugated antibodies specific for surface markers such as CD69 and PD-1, followed by washing. They were then analyzed by flow cytometry through the appropriate analysis workflow (**Figure S40**).

Tumor Immune Microenvironment (TIME) reprogramming was assessed as follows: Tumors were collected at the endpoint of the *in vivo* therapeutic efficacy study (3 weeks post-injection) and minced into ∼1–2 mm pieces. The diced tumors were enzymatically dissociated using a disaggregation buffer containing collagenase type IV and DNase I in PBS, followed by incubation at 37 °C. The resulting cell suspensions were filtered through a 40 µm cell strainer and centrifuged to collect dissociated cells, which were then resuspended in fresh PBS. Cells were stained with fluorochrome-conjugated antibodies against immune cell markers, including CD3, NK1.1, CD4, CD25, CD127, CD8, and PD-1, followed by washing. They were then analyzed by flow cytometry through the appropriate analysis workflow (**Figure S41**).

Data was analyzed using an appropriate analysis workflow in FlowJo software. Cell populations were quantified as the percentage (%) of cells exhibiting antibody-derived fluorescence intensity (FI), and detailed FI gating and sorting criteria are provided in the Supporting Information.

### 2.7. Western blot analysis

sEVs or CD8□ T cells transfected with siRNA were lysed with 1× RIPA buffer, and the total protein concentration was estimated using the BCA assay. 15 μg of protein from sEV lysates or 10 μg of protein from T cell lysates were separated by sodium dodecyl sulfate-polyacrylamide gel electrophoresis (SDS-PAGE). The separated proteins were transferred to nitrocellulose membranes and treated with primary antibodies (for sEVs: anti-CD9, anti-CD63, anti-CD81, anti-MHC I, anti-MHC II, anti-CD40, anti-CD80, anti-CD86 and anti-β-actin; for T cells: PD-1 and anti-β-actin). Secondary antibodies (anti-mouse/rat/rabbit HRP-linked) were then blotted, and immunoreactive species were detected using an ECL substrate under the bioimaging system. All blot data were normalized to the expression level of β-actin for quantification.

### 2.8. Cell viability test

Purified CD8□ T cells were co-incubated with sEVs for 2 days. For positive controls, antibodies were used to activate mouse T cells as follows: The plate was coated with 5 μg/mL CD3 antibody in PBS overnight at 4 °C or for 2 hours in the incubator at 37°C with 5% CO_2_. The coating solution was discarded, and the plate was rinsed once with PBS before culturing. CD28 antibody was added to the T cell culture medium at a concentration of 5 μg/mL. The MTT (3-[4,5-dimethylthiazol-2-yl]-2,5-diphenyl tetrazolium bromide) assay was conducted as follows: MC38-OVA cells were seeded into transparent 96-well plates at a density of 1 × 10□ cells per well. After incubation for 24 h at 37 °C, prepared CD8□ T cells (effectors) were added to the MC38-OVA cells (targets) at four different effector-to-target (E:T) ratios to assess concentration-dependent cytotoxicity over 24 h at 37 °C. Subsequently, the cells were gently washed with 1× PBS, and MTT solution was added to each well, followed by incubation for 2 h at 37 °C. After another gentle PBS wash, DMSO was added to dissolve the formazan crystals, and cell viability was determined by measuring absorbance at 540 nm.

#### *In Vivo* Safety Evaluation of sEVs

Total of 1.5 × 10□ sEVs, suspended in 100□µL PBS, were intravenously administered via tail-vein to male and female C57BL/6 mice in four groups: Ctrl, 3T3-sEV, imDC-sEV, and mDC-sEV. The Ctrl group received 100□µL PBS without sEVs. Mouse body weight was monitored every 2 days for 2 weeks post-injection (**Figure S28**). At 2 weeks, major organs (heart, liver, spleen, kidney, and lung) and muscle were collected, paraffin-embedded, sectioned, and subjected to histological analysis using hematoxylin and eosin (H&E) staining (**Figure S29**).

### 2.9. *Ex Vivo* Biodistribution Study of sEVs

sEVs were labeled with the lipophilic dye DiD and washed three times with PBS to remove unincorporated dye. A total of 1.5 × 10□ sEVs in 100□µL PBS were intravenously administered via tail-vein to male and female C57BL/6 mice in four groups: Ctrl, 3T3-sEV, imDC-sEV, and mDC-sEV. The Ctrl group received 100□µL PBS without sEVs. Mice were sacrificed at 2, 4, 8, and 24 hours post-injection, and major organs (heart, liver, spleen, kidney), as well as muscle and lymph nodes, were collected for *ex vivo* fluorescence imaging. Imaging analysis was performed using Aura Imaging Software v4.5 (Spectral Instruments Imaging, AZ, USA) and ImageJ. All samples were initially measured in a black 96-well plate prior to administration (**SI Table. 2**), and *ex vivo* fluorescence data were normalized to the 3T3-sEV signal for comparative quantification. Lymph nodes were further analyzed by immunohistochemistry (IHC) as follows: DiD-labeled sEVs were intravenously administered via tail vein to male and female C57BL/6 mice. After 3 hours, lymph nodes were harvested, sectioned, and stained with a FITC-conjugated CD8 antibody and DAPI. The stained lymph node sections were observed under a microscope and quantitatively analyzed using ImageJ software.

The lymph node uptake efficiency of DiD-sEVs was calculated using the following equation:

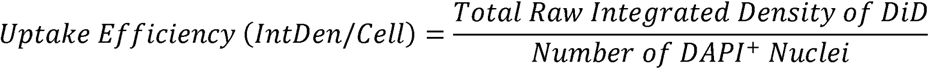

The significance of DiD-sEV colocalization within CD8[T cells was calculated as the DiD enrichment index within the FITC-positive area using the following equation:

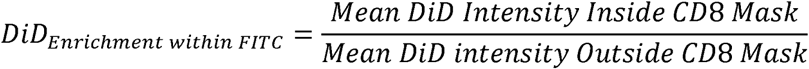

### 2.10. *In Vivo* therapeutic Study

*In vivo* therapeutic efficacy was evaluated as follows (**Figure 6a**): Mouse models were established by subcutaneously injecting 5 × 10^6^ MC38-OVA cells suspended in 50□µL of PBS into the right thigh of C57BL/6 mice. 10 days after tumor cell implantation, mice were randomly assigned to four treatment groups: Control (Ctrl), mDC-sEV, mDC-sEV(siR), and mDC-sEV(siR)/GALA. Treatments were administered intravenously via tail-vein injection. The Ctrl group received 100 µL of PBS per mouse, whereas the sEV-treated groups received 1.5 × 10^9^ sEVs per mouse in 100 µL. Treatments were administered twice, on day 0 and day 5. Tumor size was monitored on days 0, 2, 4, 7, 10, 14, 18, and 21. The representative tumor images were acquired on each measurement day. Tumor volumes were calculated using the formula:[25]

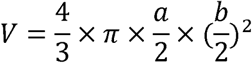

a and b represent the maximum and minimum tumor diameters, respectively. Tumor growth inhibition (TGI) was calculated using the following formula:[25]

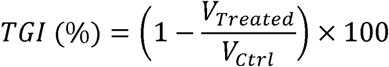

After the treatment period, major organs (heart, liver, spleen, kidney, and lung), muscle, and tumor tissues were collected, paraffin-embedded, sectioned, and stained with H&E for histological imaging analysis.

### 2.11. Statistical analysis

All data are presented as mean ± standard deviation (s.d.) or mean ± standard error (s.e.) from at least three independent experimental replicates. Statistical analysis was performed using one-way analysis of variance (ANOVA) followed by Tukey’s post hoc test. Data analysis was conducted using Microsoft Excel and SigmaPlot version 10.0. ns: not significant; *p < 0.05; **p < 0.01; ***p < 0.001; ****p < 0.0001.

## 3. RESULTS AND DISCUSSION

### 3.1. Dual-Engineered DC-sEVs Enables Efficient siRNA Loading and Cytosolic Delivery

DC maturation with Ovalbumin (OVA) was performed following established protocols[15] (see **SI Methods and Results**) and validated by robust antigen-specific activation and proliferation of OT-1 CD8□ T-cell (**Figure S1-4**). sEVs isolated from mature and immature DCs (mDC-sEVs and imDC-sEVs) showed preserved vesicle integrity and morphology by nanoparticle tracking analysis (NTA) and transmission electron microscopy (TEM) (**Figure 2a–f**), with mDC-sEVs exhibiting expected size and yield changes.[21,22,26] Functionally, mDC-sEVs induced robust OT-1 CD8□ T-cell proliferation comparable to bead-based activation, whereas imDC-sEVs showed reduced activity and non-immune mouse fibroblast-derived sEVs (3T3-sEVs) controls were inactive (**Figure 1g-h**). Compared with imDC-sEVs and 3T3-sEVs (**Figure S5**), mDC-sEVs were enriched in antigen-presentation and co-stimulatory machinery, including elevated MHC I/II, CD40, CD80, and CD86, and enhanced OVA–MHC presentation (**Figure 2i–n; Figure S6–8**).[19–21]

**Figure 2.**
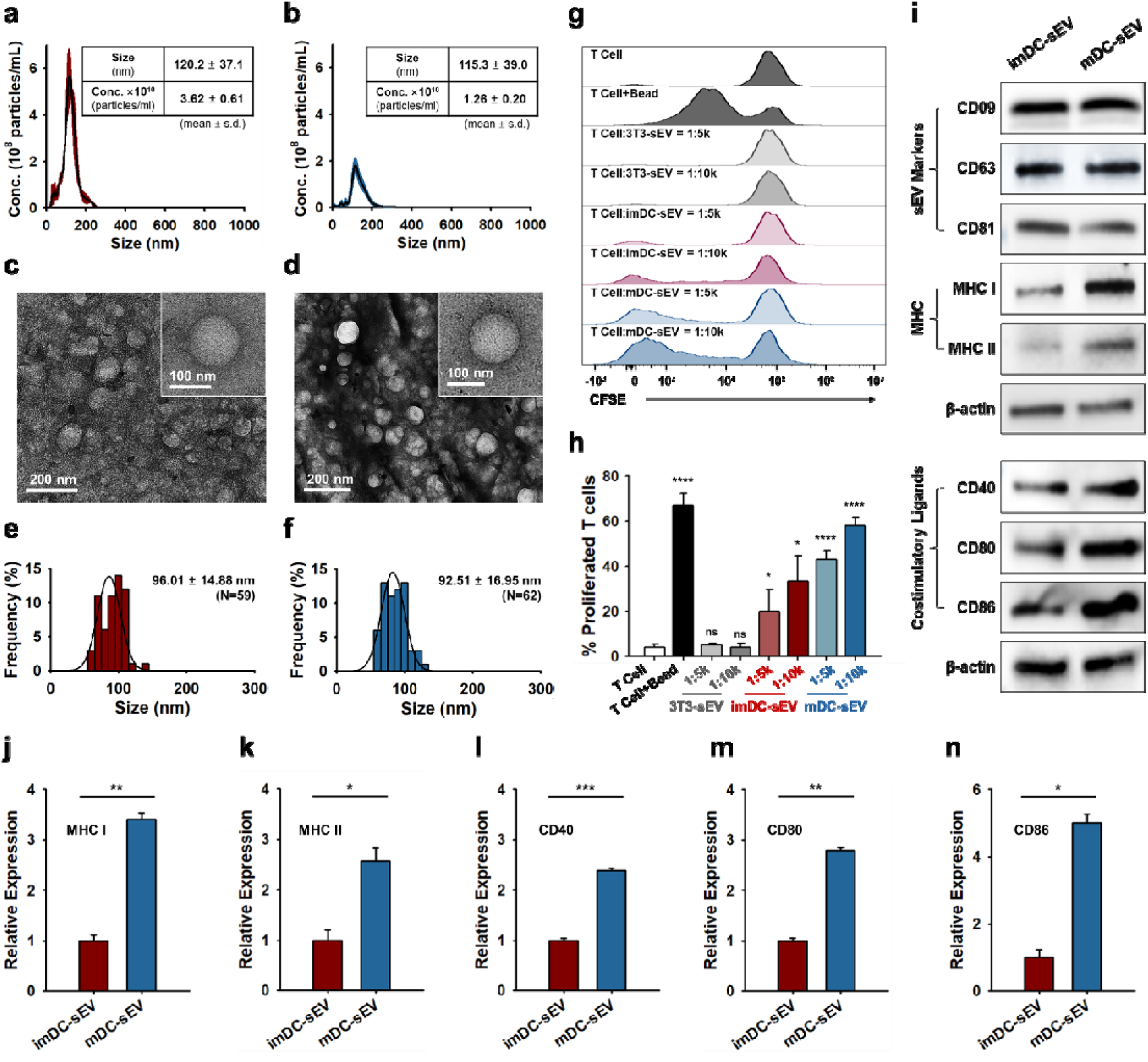
Characterization of dendritic cells derived sEVs (DC-sEVs). Size distribution and particle concentration of sEVs analyzed by nanoparticle tracking analysis (NTA) for (**a**) immature DC-sEVs (imDC-sEVs) and (**b**) mature DC-sEVs (mDC-sEVs) (n = 5, mean ± s.d). Spherical nanostructure analysis by transmission electron microscopy (TEM) and corresponding size distribution for (**c** and **e**), imDC-sEVs; (**d**) and (**f**) mDC-sEVs (mean ± s.d.). (**g** and **h**) sEV type- and dose-dependent T cell proliferation efficacy test. (**i**) Western blot analysis of sEV markers, major histocompatibility complexes (MHCs), and costimulatory ligands involved in T cell activation. Comparative expression levels of (**j**) MHC I; (**k**) MHC II; (**l**) CD40; (**m**) CD80; and (**n**) CD86 quantified from Western blot bands (n = 3, mean ± s.d).

High-efficiency siRNA loading into sEVs was achieved using a previously established chiral-assisted strategy with *D*-cysteine–functionalized graphene quantum dots (*D*-GQDs; see **SI Methods and Results**, **Figure S9**),[22] which preserve sEV integrity and enable intrinsic fluorescent cargo tracking.[9,24] GALA was incorporated via cholesterol anchoring (GALA-chol) following established protocols without compromising vesicle structure (see **SI Methods and Results, Figure S10-12**).[24] Confocal laser scanning microscopy (CLSM) imaging confirmed successful *D*-GQD loading and membrane localization before and after GALA functionalization (**Figure 3a** and **Figure S13**), with no significant change in *D*-GQD permeation efficiency (80.64 ± 5.06% vs 78.37 ± 8.34%; **Figure 3b**), indicating preserved siRNA loading capacity.[22,24] Vesicle integrity and stability were maintained, as confirmed by TEM, NTA, and time-dependent hydrodynamic measurements (**Figure 3c, d** and **Figure S14**).[22] GALA functionalization induced pH-responsive surface charge conversion (**Figure 3e**) while maintaining stability under lysosomal conditions (pH 4–5; **Figure S15**), supporting efficient endosomal escape. Flow cytometry analysis revealed GALA-density–dependent enhancement of sEV uptake by CD8□ T cells, with optimal uptake below 8 μM GALA-chol per 1 × 10□ sEVs (**Figure 3f, g**), whereas higher densities reduced uptake, likely due to steric hindrance.[27] Uptake was energy dependent, as internalization was markedly reduced at 4 °C (**Figure 3h, i**). At 37 °C, mDC-sEVs showed greater uptake than 3T3-sEVs (5.69- fold), which was further enhanced by GALA functionalization (7.92-fold). Time-course studies demonstrated maximal uptake at 6 h for mDC-sEV/GALA, exceeding 3T3-sEVs (7.92-fold), imDC-sEVs (2.67-fold), and unmodified mDC-sEVs (1.39-fold) (**Figure 3j, k**). The subsequent decline in intracellular *D*-GQD signal reflects fluorescence redistribution associated with cytosolic release rather than cargo degradation, consistent with prior reports.[24,28] These results confirm that dual engineering strategy preserves intrinsic mDC-sEV bioactivity while significantly enhancing siRNA loading and cytoplasmic delivery to T cells.

**Figure 3.**
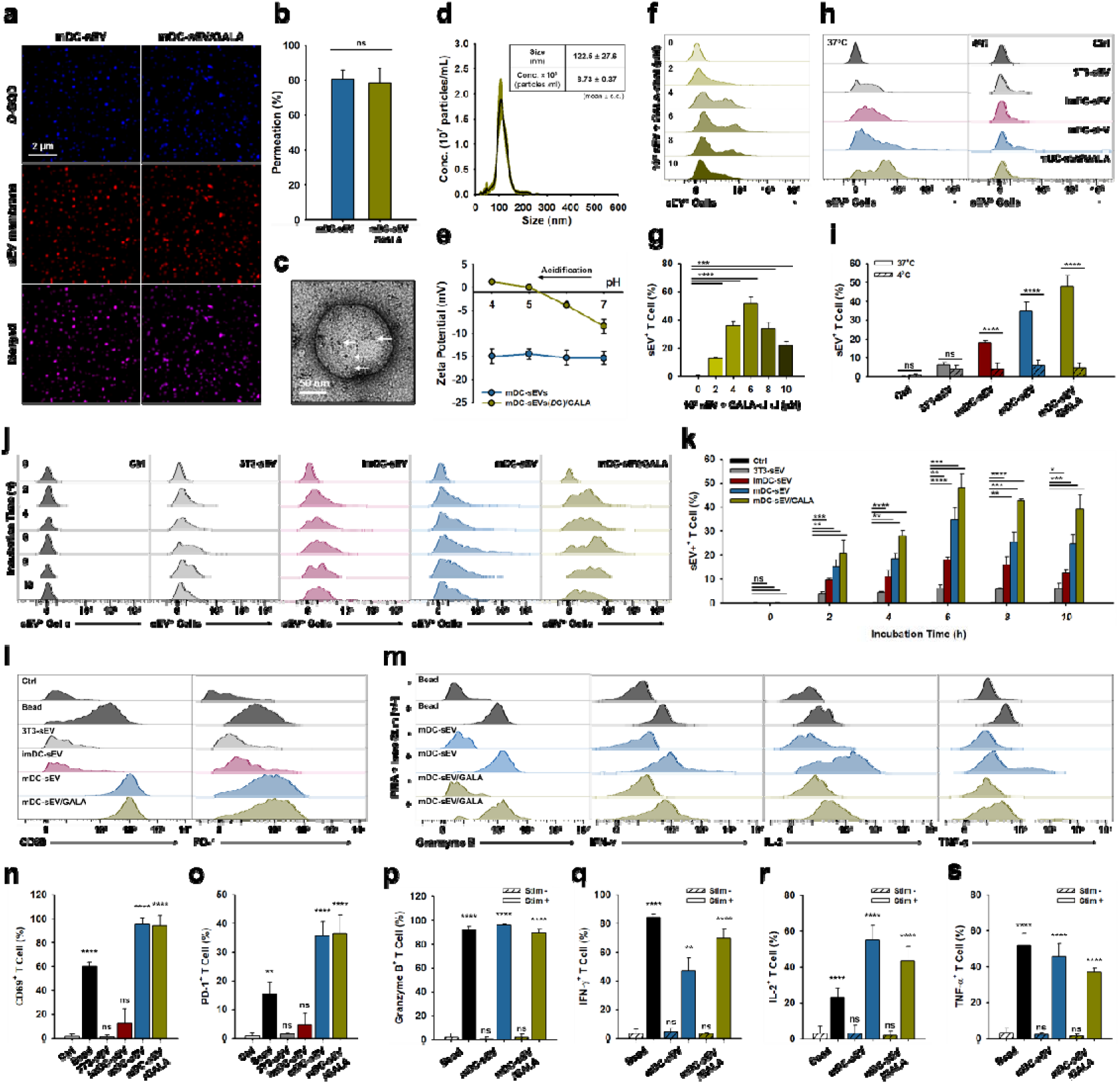
Engineering of mDC-sEV(*D-*GQD)/GALA and characterization of efficacy for T-cell activation. (**a**) Confocal laser scanning microscopy (CLSM) images of engineered mDC-sEV(*D-*GQD)/GALA. (**b**) CLSM-based quantification of *D*-GQD permeability efficiency before and after GALA functionalization (n = 4, mean ± s.d.). Each channel represents: blue for *D*-GQD; red for sEV membrane. Structural integrity preservation of mDC-sEV(*D*-GQD)/GALA analyzed by (**c**) TEM and (**d**) NTA. Arrows: *D*-GQDs within GALA-sEVs. (**e**) Surface charge change upon acidification (pH7 to pH4), corresponding to lysosomal entrapment of sEVs. (**f**) and (**g**) GALA concentration–dependent sEV uptake efficiency (GALA concentration varied on 10^9^ sEVs) after 4 h of co-incubation with OT-1 mouse–derived CD8L T cells, analyzed by flow cytometry (n = 3, mean ± s.d.). (**h** and **i**) Temperature-dependent sEV uptake efficiency after 4 h of co-incubation with OT-1 mouse–derived CD8L T cells analyzed by flow cytometry (n = 3, mean ± s.d.). (**j** and **k**) Time- and sEV type–dependent uptake efficiency analyzed by flow cytometry (n = 3, mean ± s.d.). (**l** and **n-o**) Assessment of T-cell activation analyzed by flow cytometry (n = 4, mean ± s.d.). (**m** and **p–s**) Assessment of activated T cell–mediated intracellular cytokine production analyzed by flow cytometry (n = 3, mean ± s.d.).

### 3.2. Engineered DC-sEVs Potently Activate Functional CD8□ T Cell

To confirm activation-driven proliferation and test whether GALA preserves mDC-sEV immunostimulatory function, we quantified CD8□ T-cell activation markers (**Figure 3l** and **Figure S18a, b**). Both mDC-sEV and mDC-sEV/GALA induced near-maximal CD8□/CD69□ activation (95.15 ± 5.54% and 93.86 ± 8.67%), exceeding bead activation (60.07 ± 3.67%) (**Figure 3n**). CD8□/PD-1□ populations were also higher in mDC-sEV (35.57 ± 5.00%) and mDC-sEV/GALA (36.40 ± 6.48%) than beads (15.37 ± 4.01%) (**Figure 3o**), supporting strong activation but also motivating PD-1 knockdown to mitigate PD-L1–driven dysfunction.[29]

We next assessed whether these activated CD8□ T cells were functionally competent by evaluating intracellular effector cytokine production following PMA/ionomycin restimulation (**Figure 3m** and **Figure S18c-f**). All groups produced granzyme B, IFN-γ, IL-2, and TNF-α, confirming preserved effector competence.[30] Granzyme B expression exceeded 90% in all groups, indicating strong and comparable cytotoxic potential (**Figure 3p**). IFN-γ was highest with beads (84.23 ± 2.20%), followed by mDC-sEV/GALA (69.70 ± 6.65%) and mDC-sEV (46.70 ± 9.63%) (**Figure 3q**),[31] consistent with stronger direct TCR engagement by beads[32] and enhanced cross-presentation with GALA. GALA does not act as a TCR agonist but likely increases IFN-γ by promoting lysosomal escape and cytosolic antigen delivery, enhancing cross-presentation and sustaining TCR signaling in mDC-sEV/GALA versus mDC-sEV.[20,33–36] IL-2 expression was the highest in mDC-sEVs (55.37 ± 7.69%) and modestly reduced following GALA modification (43.23 ± 8.53%), reflecting partial shielding of surface ligands despite optimized GALA density (**Figure 3r**). In contrast, bead-based activation induced substantially lower IL-2 production (23.00 ± 5.00%), consistent with its reliance on TCR-only stimulation.[37] TNF-α expression was the highest in the bead-based group (51.93 ± 6.67%), followed by mDC-sEV (45.83 ± 7.05%) and mDC-sEV/GALA (36.93 ± 2.44%), consistent with differences in the mode and strength of TCR engagement (**Figure 3s**).[30,38] Collectively, these results demonstrate that engineered mDC-sEVs elicit potent CD8□ T-cell activation while preserving broad effector functionality, and that GALA modification enhances intracellular antigen processing without compromising immunostimulatory capacity, providing a strong foundation for downstream PD-1–targeted immunomodulation.

### 3.3. Engineered DC-sEVs Enable Functional Checkpoint Blockade in CD8□ T Cells

We engineer mDC-sEVs to deliver PD-1 targeting siRNA to T cells by combining chirality–assisted *D*-GQD loading with GALA surface functionalization. After optimizing the siRNA/*D*-GQD formulation for maximal encapsulation (see **SI Methods and Results, Figure S19**), we benchmarked this passive loading approach against sonication and direct transfection. CLSM quantification[22] showed that *D*-GQD loading achieved high siRNA permeation (63.92 ± 4.54%), 10.27-fold increase over sonication (6.22 ± 1.65%) (**Figure 4a-c** and **Figure S20**). Direct transfection instead increased apparent particle size (>∼2×) and yielded fewer, partially non-colocalized Cy3-siRNA puncta, consistent with aggregation/hybrid complex formation.[9,39] In contrast, *D*-GQD loading preserved sEV integrity (**Figure 4d**) and Single-molecule localization microscopy (SMLM) confirmed robust co-localization of DiD-sEVs, Cy3-siRNA, and *D*-GQDs with strong siRNA/*D*-GQD overlap (**Figure S21**). An orthogonal agarose gel assay further verified high encapsulation efficiency (64.93 ± 8.38%), 11.05-fold higher than sonication (5.88 ± 1.79%) (**Figure 4e–g** and **Figure S22**). Loaded siRNA remained stable in FBS for 6 h, and GALA extended stability to 9 h (**Figure 4h, i** and **Figure S23**), consistent with cholesterol-enhanced membrane packing and nuclease resistance.[40,41]

**Figure 4.**
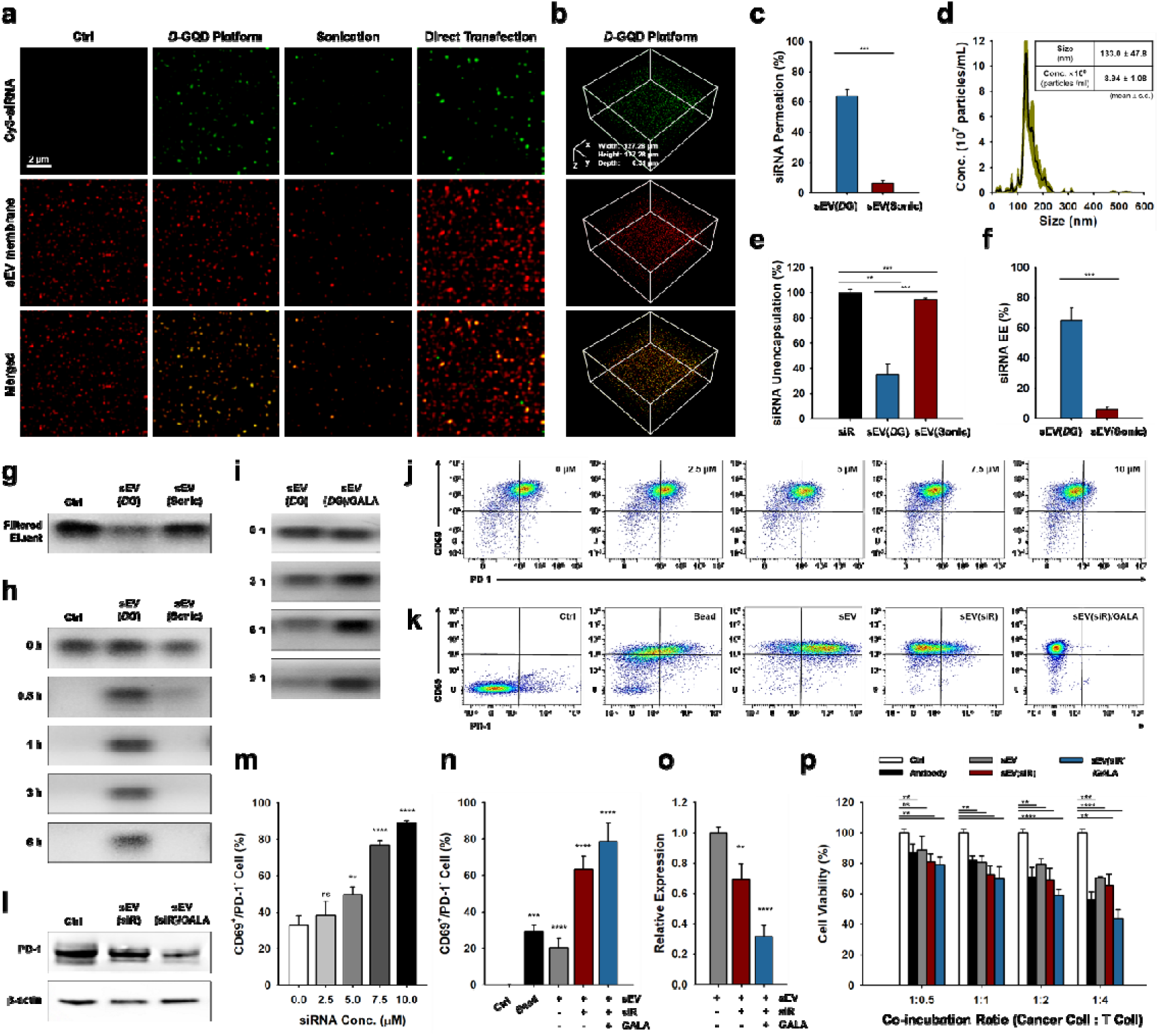
Characterization of mDC-sEV(siR)/GALA and evaluation of PD-1 knockdown efficacy. (**a**) Confocal laser scanning microscopy (CLSM) images of mDC-derived sEVs loaded with siRNA (siR). siRNA loading approach comparison: via *D*-GQD chiral-assisted loading; sonication; or direct transfection using Lipofectamine. (**b**) Z-stack CLSM images of siR-loaded sEVs prepared using the *D*-GQD loading platform. Each CLSM channel represents: green for Cy3-siRNA and red for sEV membrane. (**c**) CLSM-based quantification comparing siR permeability efficiency (n = 4, mean ± s.d.). (**d**) Structural integrity preservation of mDC-sEV(siR)/GALA analyzed by NTA. (**e–g**) Agarose gel electrophoresis analysis of washed-out eluents following siRNA loading into sEVs and corresponding quantification of siR unincorporated efficiency, followed by reverse estimation of siR encapsulation efficiency (EE; n = 3, mean ± s.d.). Agarose gel analysis of siR-loaded sEVs after incubation with fetal bovine serum (FBS), showing (**h**) time-dependent and (**i**) GALA-dependent siRNA integrity. (**j** and **m**) Dose-dependent PD-1 silencing efficacy test of mDC-sEV(siR) treated CD8L T cells analyzed by flow cytometry (n = 4, mean ± s.d.). (**k** and **n**) Comparative PD-1 silencing efficacy test of mDC-sEV treated CD8L T cells analyzed by flow cytometry (7.5 μM siR per 1 × 10^6^ CD8L T cells; n = 4, mean ± s.d.). (**l** and **o**) Western blot analysis of PD-1 expression in OT-1 mouse–derived CD8L T cells and corresponding relative expression level quantification (n = 3, mean ± s.d.). (**p**) Cancer cell viability test following co-culture with activated T cells treated with Ctrl, Antibody, sEV, sEV(siR), or sEV(siR)/GALA. Cancer cell: MC38-OVA cell line; sEV type: mDC-sEV (n = 3, mean ± s.d.).

Functionally, mDC-sEV/GALA delivered siPD-1 to CD8□ T cells and enabled checkpoint silencing. Flow cytometry showed dose-dependent PD-1 knockdown (**Figure 4j** and **Figure S24**), increasing the activated PD-1–silenced subset (CD8□/CD69□/PD-1□) (**Figure 4m**). Compared with beads and unloaded mDC-sEVs, siRNA-loaded mDC-sEVs increased this population by 3.12-fold, and GALA further boosted it to 3.87-fold (**Figure 4k, n** and **Figure S25**). Western blotting corroborated protein-level suppression (**Figure 4l, o** and **Figure S26**), with PD-1 reduced by 30.78 ± 10.42% for siRNA-loaded mDC-sEVs and by 68.32 ± 7.25% for mDC-sEV(siR)/GALA. Consistent with dual activation and checkpoint blockade, mDC-sEV(siR)/GALA–reprogrammed T cells enhanced tumor cell killing (**Figure 4p** and **Figure S27**): at a 1:4 cancer:T-cell ratio, viability decreased to 44.05 ± 5.72%, outperforming antibody-activated T cells (56.29 ± 4.95%). In sum, mDC-sEV(siR)/GALA integrates high-efficiency, serum-stable siRNA encapsulation with enhanced cytosolic delivery to achieve robust PD-1 silencing alongside strong CD8□ T-cell activation, resulting in superior functional antitumor responses.

### 3.4. Systemic Biodistribution Reveals Preserved and Prolonged Lymph Node Targeting by Engineered mDC-sEVs

To determine how parental cell origin and surface engineering affect *in vivo* trafficking, we performed time-resolved *ex vivo* fluorescence imaging of major organs (heart, liver, spleen, and kidney), lymph nodes (LN), and muscle (background) after tail-vein injection of DiD-labeled sEVs at 2, 4, 8, and 24 h (**Figure 5a–j** and **Figure S30**). All DC-sEVs showed significantly lower liver retention than 3T3-sEVs, consistent with origin-dependent systemic recognition and clearance,[42] whereas 3T3-sEVs exhibited the highest hepatic accumulation (**Figure 5k, n**). GALA functionalization modestly increased liver signal relative to control mDC-sEVs but remained far below 3T3-sEVs, suggesting enhanced reticuloendothelial system (RES)/mononuclear phagocyte system (MPS) sequestration.[43] In the spleen, DC-sEVs showed transient enrichment, peaking at 4 h (**Figure 5l, o**) and declining thereafter, consistent with rapid engagement with immune populations and subsequent trafficking rather than passive clearance.[44,45] In contrast, 3T3-sEVs progressively accumulated over time, indicative of delayed processing and prolonged sequestration.[43]

**Figure 5.**
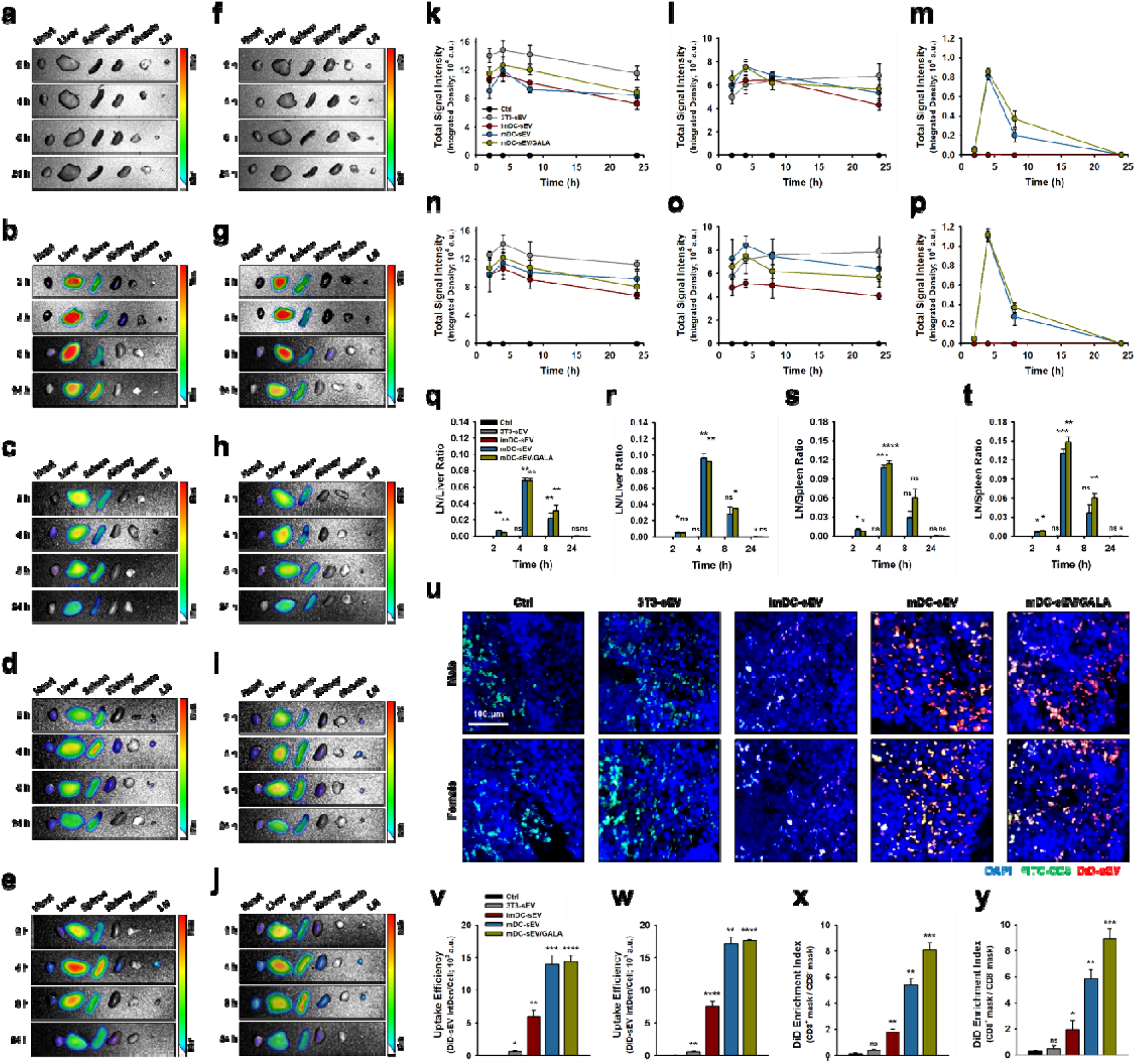
*Ex vivo* biodistribution of DiD-labeled sEVs following tail-vein i.v. injection. Representative *ex vivo* fluorescence images of major organs and lymph node (LN) from male mice: (**a**) Ctrl (PBS); (**b**) 3T3-sEVs; c, imDC-sEVs; (**d**) mDC-sEVs; and (**e**) mDC-sEVs/GALA; and from female mice: (**f**) Ctrl; (**g**) 3T3-sEVs; (**h**) imDC-sEVs; (**i**) mDC-sEVs; and (**j**) mDC-sEVs/GALA. Time-dependent quantification of sEV accumulation in major immune organs from male mice: (**k**) liver; (**l**) spleen; and (**m**) LN; and from female mice: (**n**) liver; (**o**) spleen; and (**p**) LN. LN targeting efficiency relative to liver and spleen for male mice: (**q** and **s**); and for female mice: (**r** and **t**) (n = 3, mean ± s.d.). (**u**) Immunofluorescence (IF) images of LN collected 3 h after tail-vein i.v. injection of sEVs in male and female mice. IF-based quantification of sEV uptake efficiency in LNs and sEV enrichment within CD8L T cells in male mice: (**v** and **x**); and in female mice: (**w** and **y**) (n = 3, mean ± s.d.).

DC-sEVs displayed pronounced LN accumulation, with higher overall signal in females than males (**Figure 5m, p**). Given that LNs are immune-filtering and lymphocyte-rich sites,[15] strong LN enrichment in mDC-sEV groups aligns with intrinsic immunological targeting and our *in vitro* observations. Importantly, GALA functionalization preserved LN targeting and prolonged LN-associated fluorescence versus control mDC-sEVs, consistent with delayed intracellular processing. This durability agrees with improved membrane stability in the agarose gel assay (**Figure 4i**) and may support sustained antigen presentation associated with elevated IFN-γ (**Figure 3q**). To quantify LN targeting relative to dominant off-target organs, we calculated LN/liver (**Figure 5q, r**) and LN/spleen ratios (**Figure 5s, t**): mDC-sEV/GALA almost matched mDC-sEV at 4 h and exceeded it at 8 h in both male (1.43-fold LN/liver, 2.04-fold LN/spleen) and female (1.25-fold LN/liver, 1.22-fold LN/spleen) mice.

For spatial validation, LNs harvested at 3 h were analyzed by immunofluorescence histology (**Figure 5u** and **Figure S31**). After normalizing DiD signal to cell number (see **Methods**), LN uptake of mDC-sEV and mDC-sEV/GALA was comparable and higher than imDC-sEV (2.34-fold in males, 2.41-fold in females) and 3T3-sEV (22.58-fold in males, 30.12-fold in females) (**Figure 5v, w**). To assess preferential localization within CD8□ T cell–rich regions, we quantified a DiD-sEV enrichment index within the CD8□ mask (see Methods and Figure 5x, y). Although overall uptake was similar between mDC-sEV and mDC-sEV/GALA, GALA increased CD8LJ-region enrichment (male: 1.50-fold vs. mDC-sEV; female: 1.52-fold vs. mDC-sEV). Together, these data indicate that mDC-sEV/GALA retains intrinsic LN tropism to lymphocyte-rich niches critical for antigen presentation and T cell activation[15] while enhancing persistence within CD8LJ T cell–dense regions, supporting efficient cytosolic delivery and downstream T cell reprogramming.

### 3.5. Dual-Engineered DC-sEVs Drive Antitumor Efficacy via *In Situ* T-Cell Reprogramming and Tumor Immune Microenvironment (TIME) Remodeling

We evaluated *in situ* T-cell reprogramming and the antitumor efficacy of engineered mDC-sEVs in a syngeneic MC38 murine colon carcinoma model expressing ovalbumin (MC38-OVA). MC38-OVA cells were inoculated subcutaneously into the left thigh of C57BL/6 mice. Mice were randomized into four groups: Ctrl, mDC-sEV, mDC-sEV(siR), and mDC-sEV(siR)/GALA. Starting on day 10 post-implantation, mice received two intravenous doses (tail-vein) at a 5-day interval (**Figure 6a**). Tumor growth was comparable across groups for the first 4 days, but treatment-dependent divergence emerged from day 7 (**Figure 6b, c** and **Figure S32**). In male mice, unmodified mDC-sEVs induced only a transient growth delay around day 7 and ultimately failed to improve outcome versus control (final tumor growth inhibition [TGI] vs. Ctrl: 5.12 ± 12.06%). This limited efficacy is consistent with activation-associated exhaustion, as mDC-sEV treatment increased both activation and exhaustion markers, including PD-1 (**Figure 3l** and **Figure 4l**). In contrast, siRNA delivery substantially improved anti-tumor effect. Both mDC-sEV(siR) and mDC-sEV(siR)/GALA significantly delayed tumor growth, particularly from day 14. Importantly, tumors in the mDC-sEV(siR) group began to regrow at later time points (day 21), whereas mDC-sEV(siR)/GALA maintained suppression through study end (final TGI vs. Ctrl: 61.80 ± 7.60% and 82.12 ± 4.49%). These results indicate enhanced intracellular siRNA delivery and PD-1 silencing of engineered mDC-sEVs *in vivo*, sustaining T-cell effector function and limiting exhaustion.

**Figure 6.**
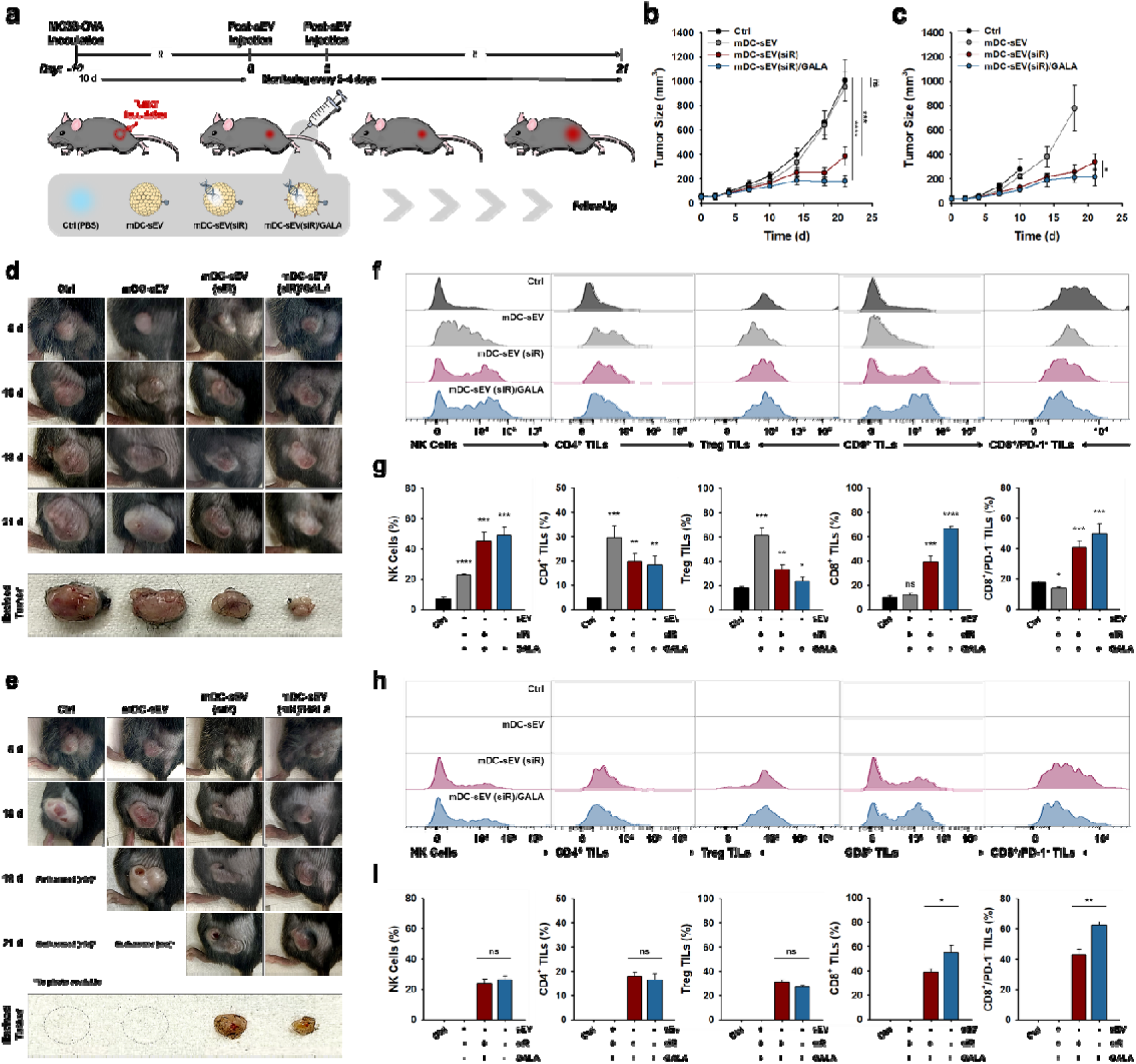
*In vivo* therapeutic efficacy via *in situ* T-cell reprogramming following systemic administration of mDC-sEV(siR)/GALA. (**a**) Schematic workflow illustrating tumor progression monitoring after tail-vein i.v. administration of sEV treatments. Tumor volume growth curves in (**b**) male mice, and (**c**) female mice following treatment (n = 4, mean ± s.d.). Representative photographs of MC38-OVA tumor–bearing mice taken at 0, 10, 18, and 21 days post-injection in (**d**) male mice, and (**e**) female mice. Assessment of tumor immune microenvironment (TIME) reprogramming following mDC-sEV treatment analyzed by flow cytometry. Immune cell populations in dissociated tumors collected at the termination of the *in vivo* therapeutic efficacy study (day 21 post-injection). (**f** and **g**) Immune cell population analysis in male mice; and (**h** and **i**) in female mice. Populations compared include: natural killer (NK) cells, CD4^+^ tumor-infiltrating lymphocytes (TILs), regulatory T cells (Treg TILs), CD8^+^ TILs, and CD8^+^/PD-1^-^ TILs (n = 3, mean ± s.d.).

Sex-dependent immune differences have been reported in syngeneic tumor models [46,47] and female mice often exhibit stronger antigen-specific T-cell responses,[47,48] which can accelerate immune-mediated tumor killing and necrosis in highly immunogenic MC38-OVA tumors.[49] Consistently, female tumors showed increased morbidity (Ctrl euthanized by day 10; mDC-sEV: 50% survived beyond day 10 [n=2] but not to day 21). Nevertheless, both siRNA–loaded groups maintained all surviving replicates to study end, and siRNA delivery delayed progression with the most sustained control in the GALA-functionalized group (final tumor volumes: 339.44 ± 65.78 mm³ for mDC-sEV(siR) and 213.62 ± 70.69 mm³ for mDC-sEV(siR)/GALA). Endpoint tumors from siRNA-treated mice displayed irregular, heterogeneous architectures in males (**Figure 6d**) and females (**Figure 6e**), consistent with localized T-cell reactivation following PD-1 silencing and focal regression, vascular disruption, and fibrotic remodeling,[50,51] accompanied by elevated tumor-infiltrating lymphocytes (TILs) and immune-associated apoptosis (**Figure S33, 34**). Given that low immune infiltration and immunosuppression enable tumor escape,[15,52] we profiled tumor immune microenvironment (TIME) remodeling at endpoint by flow cytometry (**Figure 6f–i** and **Figure S35**). NK cell infiltration increased in both siRNA-delivering groups (higher in males). CD4□ T cells expanded most in the unloaded-mDC-sEV group, which also increased intratumoral Tregs, suggesting a shift toward regulatory subsets. In contrast, CD8□ T-cell infiltration tracked tumor control: mDC-sEV(siR)/GALA produced the highest CD8□ T-cell levels, and PD-1–silenced CD8□ T cells were enriched in siRNA-loaded groups, with the highest abundance in the GALA group, consistent with its durable suppression (**Figure 4k**). These data support that mDC-sEV alone elicits activation coupled to PD-1 upregulation, limiting durability, whereas PD-1 silencing enhanced by GALA reprograms the TIME toward cytotoxic immunity, paralleling principles of DC-sEV–mediated TIME remodeling[15] and ICB effect.[53–55]

### 3.6. Discussion

In contrast to current cell-based immunotherapies (e.g., CAR-T cells and DC vaccines), which require extensive *ex vivo* manipulation and reinfusion and thus face barriers in complexity, cost, scalability, and variability,[1,56] mDC-sEVs drive *in situ* T-cell reprogramming without viable cell transfer. Moreover, unlike systemic ICB that broadly circulates and can elicit off-target immune effects,[3] mDC-sEVs consolidate antigen presentation, co-stimulation, and localized ICB cargo delivery to control activation at the tumor–immune interface, supporting off-the-shelf manufacturing and storage.[15,57] Unlike CAR-T cells with fixed specificity and persistence-linked toxicities,[58] and DC vaccines with inefficient trafficking and short *in vivo* lifespan,[59] mDC-sEVs preserve DC immunobiology that supports lymphoid targeting (**Figure 5k–p**), while surface engineering further enhances LN-normalized persistence (**Figure 5q–t**), enabling sustained yet programmable activation without permanent genetic modification.[15] Notably, because solid-tumor efficacy is frequently blunted by an immunosuppressive TME, mDC-sEVs are designed to function within the TIME and shift immune composition—including NK cells, CD4□/CD8□ T cells, and Tregs—toward an effector-supportive, tumor-specific response (**Figure 6f–i**).[60–62]

DCs are compelling sEV sources for cancer immunotherapy given their role as professional antigen-presenting cells that prime T-cell responses.[14,15] Early studies[63] demonstrated that antigen-equipped DC-sEVs can induce antitumor immunity and improve outcomes in murine melanoma[64] and lung carcinoma models.[65] However, DC-sEV–driven T-cell activation alone is often insufficient because tumors impose immune evasion and checkpoint–mediated T-cell exhaustion.[2,29] Effective immunity therefore benefits from integration of ICB strategies,[3] but sEV delivery of intracellular ICB agents remains limited by inefficient/stochastic loading,[66,67] and lysosomal sequestration that restricts cytosolic bioavailability.[68,69] Our dual-engineering strategy addresses these barriers by combining efficient ICB cargo loading and lysosomal escape with preserved DC-sEV bioactivity, enabling sustained T-cell–mediated efficacy.

Finally, building on DC-to-sEV transfer of exogenously processed peptide–MHC complexes, we anticipate that “antigen-educating” DCs (e.g., with tumor lysates or patient-derived antigens) could tailor mDC-sEV specificity for personalized T-cell priming across cancers,[15] while rational antigen selection and added immunomodulatory cues may extend the platform toward cancer type–agnostic therapeutics or vaccines.

## 4. CONCLUSION

In this study, we established a dual-engineered DC-sEV platform for localized *in situ* T-cell reprogramming that couples preserved DC immunobiology with programmable genetic checkpoint modulation. By retaining DC-derived antigen-presentation and costimulatory machinery while enabling high-efficiency siPD-1 loading and cytosolic delivery, engineered mDC-sEVs integrate endogenous immune signaling with targeted ICB gene silencing to drive robust antitumor activity and remodel the TIME, sustaining effector function and coordinated antitumor immunity.

## ACKNOWLEDGEMENTS

TEM, CLSM, *ex vivo*, and H&E histology images were obtained using the instruments of Notre Dame Integrated Imaging Facility (NDIIF), University of Notre Dame. NTA analysis was obtained using the instrument of Harper Cancer Research Institute (HCRI), University of Notre Dame. FTIR analysis was obtained using the instrument at the Center for Environmental Science and Technology (CEST), University of Notre Dame. Western blot and agarose gel images were acquired using the instrument of Biophysics Instrumentation Core (BIC), University of Notre Dame. All animal studies were conducted at Freimann Life Science Center (FLSC), University of Notre Dame. We appreciate the support from these facilities for this study. The authors gratefully acknowledge Dr. Sihan Yu and Dr. Bowen Fan for their valuable contributions to the development of the modified GALA peptides. This study was supported in part by the Berry Family Foundation Graduate Fellowship from the Berthiaume Institute for Precision Health (BIPH; to GK) at the University of Notre Dame. This study was supported in part by the Interdisciplinary Interface Training Program (IITP) Grant from HCRI, with funding from the Walther Cancer Foundation (WCF; to GK and SW). This work was supported by National Institutes of Health (NIH) under the Maximizing Investigators’ Research Award (MIRA) [R35GM150608; YW] and National Institutes of Health Grant [R21CA277663-01A1; YW]. Other supports included NIH grants R01CA248033 (XL), R01CA280097 (XL), R01CA297220 (XL), and Department of Defense grants W81XWH2010332 (XL), W81XWH2010312 (XL), HT94252310010 (XL) and HT94252310613 (XL).

## AUTHOR INFORMATION

### Author Contributions

Y.W., X.L. and G.K. conceptualized the study. G.K. designed the methodology and carried out all the experiments, as well as the data analysis. S.W. isolated T cells and contributed to the flow cytometry study under the supervision of X.L. R.Z. synthesized and characterized chiral GQDs. M.J.W designed and developed the modified GALA peptides. G.K. wrote the original manuscript and created all schematic illustrations in this manuscript. Y.W. and X.L. supervised the entire project, including manuscript writing. All authors contributed to data interpretation, discussion, and writing.

## STATEMENTS AND DECLARATIONS

### Ethical Approval Statement

Animals involved in this study under protocol 22-08-7344 and 24-11-8960 that were reviewed and approved by The University of Notre Dame’s Institutional Animal Care and Use Committee (IACUC).

