## Supplementary material for "Dual-Engineered Dendritic Cell–Derived Small Extracellular Vesicles Couple T-Cell Priming with Checkpoint Reprogramming for Synergistic Immunotherapy": SI File

### SUPPLEMENTARY METHODS

#### Synthesis and Characterization of Chiral Graphene Quantum Dots (GQDs)

The GQDs were synthesized using a modified Hummers' method based on our previous studies as follows:[1,2] Carbon nanofibers were used as a precursor for top-down GQD synthesis via oxidative treatment with $H_2SO_4$ (98%) and $HNO_3$ (68%) at 120 °C for 20 h. The product was neutralized with NaOH, purified by dialysis (1 kDa MWCO), followed by surface-functionalized with *D*-cysteine via EDC/NHS coupling. Chiral GQDs were further purified by dialysis and filtered through a 0.22 µm syringe filter. Characterization included transmission electron microscopy (TEM; 200 kV), fluorescence spectroscopy (microplate reader), circular dichroism (CD) spectroscopy for optical absorbance and chiroptical properties, and Fourier-transform infrared spectroscopy (FTIR) for chemical composition.

#### Synthesis and Characterization of Cholesterol Conjugated pH-Responsive Peptides

Modified GALA peptides (GALA-chol and FAM-GALA-chol) were synthesized according to our previous study, as follows:[3] A cysteine-modified GALA peptide (Ac-WEAALAEALAEALAEHLAEALAEALEALAAC-$NH_2$) was synthesized via solid-phase methods using a CEM Liberty Blue synthesizer. The cholesterol linker was prepared by reacting cholesterol with bromoacetic acid in the presence of DIC and DMAP in anhydrous dichloromethane. Subsequently, the GALA peptide and its FAM-labeled variant were conjugated to the cholesterol linker through thioether linkage via the C-terminal cysteine residue.

#### Characterization of siRNA/*D*-GQD Complex

The siRNA/*D*-GQD complex was characterized by fluorescence change analysis:[1] A fixed concentration of Cy3-siRNA (10 µM) was incubated with increasing concentrations of *D*-GQDs (0–5 µM). *D*-GQD (Ex/Em 360/510 nm) served as the donor and Cy3–siRNA (Ex/Em 550/570 nm) as the acceptor. Fluorescence resonance energy transfer (FRET) was monitored as Cy3 emission at 570 nm upon selective donor excitation at 360 nm. FRET-induced fluorescence increased upon binding of Cy3-siRNA to *D*-GQDs, reaching a plateau at higher *D*-GQD concentrations. Relative fluorescence change due to FRET was calculated using the following equation:

$$Relative\ FRET = \frac{\mathrm{F} - F_0}{F_0}$$

The binding efficiency plot was generated from normalized values using the Relative FRET equation.

To determine the optimal incubation time for *D*-GQDs and siRNA/*D*-GQD complexes to permeate into sEVs, the following procedure was used: *D*-GQDs were incubated with sEVs for varying time points (0–20 min). Unpermeated *D*-GQDs were removed using 100 kDa centrifugal filters to isolate sEVs, which were then lysed to release internalized *D*-GQDs. Fluorescence of *D*-GQDs was measured at Ex/Em = 360/510 nm.[4] Similarly, Cy3-siRNA/*D*-GQD complexes were subjected to the same procedure, and FRET fluorescence was measured at Ex/Em = 360/570 nm.

**Characterization of siRNA Encapsulation**

The siRNA encapsulation efficiency was quantified using an agarose gel method specifically developed in our previous study:[3] The washed-out eluent containing unbound siRNA removed from the sEVs was collected after filtration. This siRNA was then run on an agarose gel electrophoresis, and the bands were analyzed to estimate the amount of unencapsulated siRNA. Subsequently, a reverse estimation was performed to quantify the amount of siRNA loading within the sEVs. Data from each round were normalized to the expression level of the naked siRNA group. The encapsulation efficiency was calculated using the following equation:

$$Unencapsuation\ (Loss;\ \%) = \frac{siRNA_{Filtrate}}{siRNA_{Free}} \times 100$$

$$Encapsulation\ Efficiency\ (EE\%) =\ 100 - Loss\ (\%)$$

For the time-dependent siRNA integrity test against RNase under physiological conditions, siRNA-loaded sEVs were incubated with 4% FBS at 37 °C for 0–9 h. After each corresponding time, the sEVs were filtered to remove FBS and treated with a mild surfactant (0.2% Triton X-100) to induce leakage and extract the siRNA. This siRNA was then run on an agarose gel electrophoresis, and the bands were analyzed to evaluate the degradation. For the agarose gel electrophoresis described above, all samples were mixed with 6× DNA loading buffer and loaded onto a 1% TBE agarose gel containing EtBr. Electrophoresis was performed at 120 V for 30 min, and siRNA was visualized using a bioimaging system with 365 nm excitation.

**Confocal Laser Scanning Microscopy (CLSM) Imaging and Analysis of sEVs**

The permeation efficiency of *D*-GQDs and/or Cy3-siRNA into sEVs was calculated using a formula from our previous report and analyzed statistically:[1] 5 μL sample of *D*-GQD–loaded sEVs or siRNA/*D*-GQD–loaded sEVs was applied onto Poly-L-Lysine–coated slides (63410; Electron Microscopy Sciences, PA, USA) and fully covered with 18 mm × 18 mm square cover glasses (2845-18; Corning, NY, USA). The covered area was scanned under a 100× objective with a field size of 127.28 μm × 127.28 μm. Each scan included 25 z-stack images (step size 0.125 μm), capturing the appearance and disappearance of blue (*D*-GQD) or green (Cy3-siRNA) fluorescent dots. Based on the captured images, the permeation efficiency was calculated using the following equation:[3]

$$Permeation\ Efficiency\ (\%) = \frac{\sum(TFEPs - CFEPs)}{4 \times TEC} \times \frac{18\ mm \times 18\ mm}{127.28\ \mu m \times 127.28\ \mu m} \times \frac{1}{5\ \mu L} \times 100$$

Successful hydrophobic insertion of GALA-chol into the sEV membrane was confirmed using the same CLSM imaging procedure, and the results were statistically quantified as follows: Pearson's correlation coefficient (PCC) was used to quantify colocalization between two channels (green: FAM-GALA-chol and red: sEV membrane). PCC values range from –1 to 1, where –1 indicates perfect negative correlation, 0 indicates no correlation, and 1 indicates perfect positive correlation. Higher PCC values reflect greater degrees of colocalization. The scatter plots for the two channels were generated using ImageJ software with the JACOPx Plugin. The PC linear plot was derived from these scatter plots, followed by calculating the PCC using the following equation:[3]

$$PCC = \frac{\sum(C1_i - C1_{av}) \times (C2_i - C2_{av})}{\sqrt{\sum(C1_i - C1_{av})^2 \times \sum(C2_i - C2_{av})^2}}$$

$C1_i$ and $C1_{av}$ denote the intensity values and the average intensity of channel 1, respectively, while $C2_i$ and $C2_{av}$ denote the intensity values and the average intensity of channel 2. Representative CLSM figures in this study were obtained from 50 z-stack acquisitions and were presented as either 2D cumulative single-plane projections or 3D cube-structured reconstructions. Furthermore, SMLM was used to image individual sEVs stained with DiD (red), loaded with siRNA (Green) and *D*-GQDs (blue) complexes.

### SUPPLEMENTARY RESULTS

#### Enhanced T Cell Stimulation by MHC-Enriched sEVs from Mature Dendritic Cells

To enrich MHC presentation on DC-derived sEVs, DCs were stimulated with ovalbumin (OVA) to induce maturation (**Supplementary Fig. 1, 2**).[5] Mature DCs were first validated functionally by co-cultured with OT-1 CD8$^+$ T cells (OVA-specific TCR; **Supplementary Fig. 3**), which confirmed that robust antigen-specific CD8$^+$ T cells activation and proliferation (**Supplementary Fig. 4**). sEVs isolated from mature DCs (mDC-sEVs) and immature, non–OVA-stimulated DCs (imDC-sEVs)[1,6] both displayed narrow size distributions by nanoparticle tracking analysis (NTA), indicating preserved vesicle integrity (**Fig. 2a, b**). Compared with imDC-sEVs, mDC-sEVs were slightly smaller and yielded ~3-fold fewer particles, consistent with phenotypic and metabolic changes during DC maturation.[7] Transmission electron microscopy (TEM) further confirmed the spherical morphology and intact structure of the DC-sEVs (**Fig. 2c–f**). Consistent with selective cargo sorting and membrane compartmentalization during sEV biogenesis that concentrates surface ligands,[7–9] mDC-sEVs exhibited significantly higher immunostimulatory ligand expression than their parental mDCs (**Supplementary Fig. 6**). Flow cytometry analysis demonstrated that mDC-sEVs were strongly enriched in antigen-presentation and co-stimulatory machinery relative to imDC-sEVs. Specifically, mDC-sEVs showed increased surface MHC I and MHC II (3.40- and 2.57-fold, respectively), upregulated co-stimulatory ligands (CD40, 2.38-fold; CD80, 2.79-fold; CD86, 5.00-fold) (**Fig. 2i–n** and **Supplementary Fig. 7**), and enhanced OVA-specific peptide–MHC presentation (**Supplementary Fig. 8**). To benchmark immune specificity, we included sEVs derived from 3T3 fibroblasts (3T3-sEVs) as a non-immune control in all comparisons. At a T cell–to–sEV ratio of 1:10,000, mDC-sEVs induced 58.17 ± 3.44% OT-1 T-cell proliferation, comparable to a commercial bead-based activation system, whereas imDC-sEVs elicited only 33.33 ± 11.45% under identical conditions (**Fig. 2h**). In contrast, 3T3-sEVs failed to stimulate T-cell proliferation and remained similar to untreated controls, consistent with minimal expression of MHC and co-stimulatory ligands (**Supplementary Fig. 5**).

#### Design and Characterization of Chiral Chiral GQDs as a Gene-Loading Platform for sEVs

Chiral GQDs were engineered as an efficient platform for siRNA loading into sEVs by functionalizing GQDs with chiral amino acids, such as cysteine, following our previously reported methods.[1] CD spectroscopy demonstrated pronounced chiroptical activity in *D*-cysteine–

modified GQDs (*D*-GQDs) (**Supplementary Fig. 9e**). In addition, *D*-GQDs exhibited a red-shifted fluorescence emission spectrum relative to pristine GQDs (**Supplementary Fig. 9f**), which is attributed to the electron-donating properties of *D*-cysteine. Consistently, UV–vis absorbance spectra also showed a corresponding shift (**Supplementary Fig. 9g**), further confirming successful chiral functionalization. The chemical composition of *D*-GQDs was characterized by FTIR spectroscopy (**Supplementary Fig. 9c**), revealing characteristic absorption bands at 1706 $cm^{-1}$ corresponding to C=O stretching and at 1250 $cm^{-1}$ associated with C–N stretching from *D*-cysteine, in contrast to unmodified GQDs. TEM analysis (**Supplementary Fig. 9a, b**) indicated an average particle diameter of 6.23 ± 2.43 nm. This size was deliberately optimized to approximate lipid bilayer thickness, facilitating efficient permeation into sEVs (**Supplementary Fig. 9d**).[4]

**pH-Responsive Surface Functionalization of sEVs Using GALA-Chol**

To enable pH-responsive lysosomal escape and enhance cytoplasmic gene delivery, sEVs were functionalized with a cysteine-modified GALA peptide. The GALA sequence (Ac-WEAALAEALAEALAEHLAEALAEALEALAAC-$NH_2$) was synthesized and site-specifically conjugated with a cholesterol moiety at the C-terminal cysteine residue, yielding GALA-chol (**Supplementary Fig. 10**).[3] The cholesterol anchor inserts into the sEV lipid bilayer via hydrophobic interactions, allowing surface presentation of the GALA peptide. This spontaneous, passive lipophilic association preserves the intrinsic membrane bioactivity of sEVs,[10] while imparting pH-responsive modulation of surface charge (**Fig. 3e**).[3] Upon acidification, the resulting surface charge modulation promotes endo-lysosomal membrane fusion between the neutrally to positively charged sEV membrane and the negatively charged lysosomal membrane, thereby facilitating efficient lysosomal escape and cytoplasmic cargo delivery.[3]

Additionally, we further confirmed the successful incorporation of the GALA peptide onto sEVs. We introduced fluorescein (FAM)–labeled GALA-cholesterol (FAM-GALA-chol) into sEVs, and fluorescence recovery was monitored following membrane lysis using the biological surfactant Tween-20. The evident fluorescence increase after lysis indicates that partial self-quenching of FAM on intact sEVs was alleviated upon membrane disruption, which increased the intermolecular distance between fluorophores and thereby enhanced fluorescence by reducing quenching effects in the lysed suspension. We first optimized the duration of the hydrophobic interaction between FAM-GALA-chol and sEVs and found that fluorescence recovery reached saturation after 10 min of co-incubation with 1 × $10^9$ mDC-sEVs at 4 μM FAM-GALA-chol, following membrane lysis with the biological surfactant Tween-20 (**Supplementary Fig. 11a, b**). Subsequently, concentration-dependent studies of FAM-GALA-chol revealed a proportional increase in recovered fluorescence intensity (**Supplementary Fig. 11c, d**). Moreover, FAM-GALA-chol–functionalized mDC-sEVs were further stained with the lipophilic dye DiD to label the sEV lipid bilayer and characterized by CLSM using both 2D (single-plane) and 3D (Z-stack) analyses (**Supplementary Fig. 12**). Colocalization analysis revealed a high Pearson's correlation coefficient (PCC = 0.947), indicating strong spatial correlation between FAM (green) and DiD (red) fluorescence signals. These results collectively demonstrated the efficient membrane incorporation of cholesterol moiety and successful surface display of the GALA peptides on sEVs.

To demonstrate the pH-responsive surface charge conversion of sEVs via cholesterol-mediated GALA peptide insertion, we analyzed the surface charge of sEVs (**Supplementary Fig. 16**) to assess whether GALA-chol efficiently integrates into the sEV membrane and enables controlled charge modulation, as measured by zeta potential analysis (**Supplementary Fig. 17a, b**). The results showed that GALA-functionalized sEVs adjusted their surface charge to neutral or slightly positive values under endo-lysosomal pH conditions (pH 4–5),[3] which is favorable for lysosomal escape and correlates with enhanced cargo uptake by T cells. Notably, at pH 3, the zeta potential profiles of sEVs, regardless GALA functionalization, exhibited highly noisy and random peak distributions (**Supplementary Fig. 17c, d**). This behavior is attributed to hydrolysis of ester linkages in phospholipid molecules under strongly acidic conditions, which compromises lipid bilayer stability and alters the structural and functional integrity of biological membranes.[3,11] Consistent with these observations, nanoparticle tracking analysis (NTA) revealed unstable particle morphology and a reduced particle concentration at pH 3 (**Supplementary Fig. 15e**), indicating membrane disruption and leakage of sEV cargo, in agreement with the zeta potential measurements.

#### Efficient siRNA Loading into sEVs via a Chiral GQD Nanoplatform

Finely size-tuned *D*-GQDs interact with siRNA through pi stacking, thereby markedly enhancing siRNA loading into sEVs via the high permeation efficiency of the siRNA/*D*-GQD complexes.[1,3] This behavior arises from selective nanoscale interactions between the twisted nanosheet architecture of *D*-GQDs and the lipid bilayers of sEVs, enabling efficient cargo translocation while preserving sEV structural integrity.[1,4] As a proof of concept, Cy3-labeled siRNA (5 μM; 21 bp) was complexed with increasing concentrations of *D*-GQDs (0–5 μM). In this FRET-based assay, *D*-GQDs (Ex/Em: 360/510 nm) served as the donor and Cy3–siRNA (Ex/Em: 550/570 nm) as the acceptor (**Supplementary Fig. 19g**). Upon complex formation, FRET-induced Cy3 fluorescence increased in a *D*-GQD concentration–dependent manner and reached saturation at higher *D*-GQD concentrations (**Supplementary Fig. 19a–d**). Quantification of the relative fluorescence change (Ex/Em: 360/570 nm) (**Supplementary Fig. 19e**), normalized to binding efficiency, revealed that each *D*-GQD was capable of carrying approximately two siRNA molecules (**Supplementary Fig. 19f**). We next optimized the complexation kinetics, demonstrating that fluorescence recovery plateaued within 10 min, with minimal standard deviation, indicating high reproducibility (**Supplementary Fig. 19h**). Subsequently, optimization of the sEV permeation duration for siRNA/*D*-GQD complexes revealed a notable reduction in variability after 10 min (**Supplementary Fig. 19i**), reflecting enhanced stability and reproducibility of the loading process. Compared with conventional loading methods, the *D*-GQD platform exhibited superior loading efficiency and permeability (**Supplementary Fig. 20**). Finally, successful encapsulation of siRNA/*D*-GQD complexes within individual mDC-sEVs was directly confirmed by single-molecule localization microscopy (SMLM) (**Supplementary Fig. 21**), validating the robustness and precision of this loading strategy.

***Ex Vivo* Biodistribution Study - Prolonged Systemic Circulation potential of mDC-SEV**

Remarkably, higher accumulation of DC-sEV signals in the heart was consistently observed in both male and female mice (**Supplementary Fig. 30**). Although these measurements were obtained from *ex vivo* imaging and not directly reflected in all blood-pool signals due to tissue dissection and blood loss during harvesting one of the key circulatory organs, elevated cardiac signals were still evident compared to non-immune cell-derived sEVs (3T3-sEVs). This may be attributed to the ability of DC-sEVs to evade immune surveillance and reduce sequestration by the reticuloendothelial/mononuclear phagocyte system (RES/MPS),[12] resulting in prolonged systemic circulation. This result is consistent with previous reports describing immune cell–derived sEVs as having enhanced immune-evasive properties,[13,14] with a more pronounced effect observed for mDC-sEVs compared with imDC-sEVs, potentially due to differences in immunomodulatory capacity, and thus, extended circulation would increase the likelihood of passive or opportunistic accumulation in highly perfused organs such as the heart. The concomitant increase in renal signal further supported delayed clearance kinetics, with kidney-associated fluorescence gradually increasing over time up to 8 h post-injection. During prolonged circulation, mDC-sEVs may be increasingly exposed to circulating enzymes or complement factors, which can partially digest their membranes and produce smaller lipid–protein complexes, which, in turn, are subsequently cleared through renal filtration.[15,16] In parallel, elevated cardiac accumulation may also be influenced by active interactions with resident cardiac immune populations, including macrophages and dendritic-like cells, which internalize DC-sEVs via receptor-mediated active uptake mechanisms,[17–19] thereby contributing to prolonged retention within cardiac tissue. Collectively, these observations suggested that the observed cardiac enrichment of mDC-sEVs is not only active cardiac targeting, but also likely arises from a combination of factors, including extended blood residence time, which confers prolonged systemic circulation—a property generally advantageous for nanoparticle-based delivery.[20] This prolonged circulation may also underlie the enhanced lymph node accumulation observed for mDC-sEVs (**Fig. 5**), supporting their favorable biodistribution profile for systemic immunotherapy.

**Histological Analysis of Major Organs and Tumor Post-Treatment**

Hematoxylin and Eosin (H&E)-stained tumor sections were analyzed at day 21 and at the endpoint of each group (with female groups having variable termination time points). No significant histopathological changes were observed in major organs (heart, lung, liver, spleen, kidney) or muscle across all groups (**Supplementary Fig. 33**). Tumors in the control group appeared fresh and well-organized, with clear contours and intact tissue architecture. In contrast, tumors from the mDC-sEV-treated group displayed heterogeneous and less organized structures with darker staining. Magnified sections (**Supplementary Fig. 34**) revealed the presence of tumor-infiltrating lymphocytes (TILs; arrows), identified as small, darkly stained, circular nuclei.[21,22] However, the abundance of TILs in the mDC-sEV groups was relatively low, likely reflecting T cell exhaustion associated with PD-1 overexpression, despite efficient T cell activation indicated by high CD69 expression *in vitro* (**Fig. 4k**). These histological observations were also consistent with tumor growth curves, which showed similar progression to the control group at the endpoint (**Fig. 6b, c**). In contrast, tumors treated with siRNA-loaded mDC-sEVs displayed clear signs of cellular stress, including abundant TILs, nuclear shrinkage, and reduced vibrancy. The extracellular matrix and cytoplasm appeared more prominent relative to nuclei, suggesting loosening of the tumor microenvironment (TME), likely driven by immune cell infiltration and activity, consistent with remodeling of the tumor immune microenvironment (TIME) population (**Fig. 6f–i**). Tumors in the sEV(siR)/GALA group exhibited similar features, but more pronounced, with occasional swirl-like, multicolored regions and disrupted nuclei, potentially indicative of TIL-mediated cancer cell attack (circles in figure).[21,22] Overall, these histological observations indicate that siRNA-loaded and GALA-functionalized sEVs promoted TME re-configuration and immune infiltration, correlating with the improved anti-tumor activity observed *in vivo* (**Fig. 6**).

### SUPPLEMENTARY FIGURES

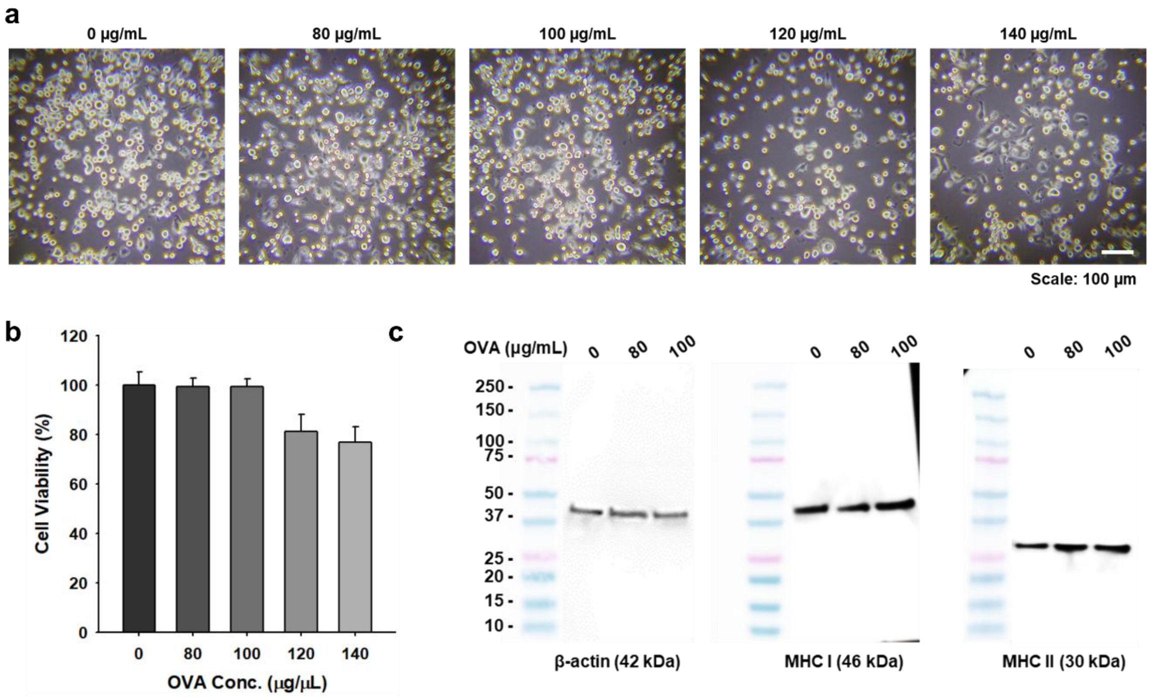


**Supplementary Fig. 1**: Dose-dependent ovalbumin (OVA)-induced maturation of dendritic cells (DCs; JAWS II cell line). (**a**) Representative microscopic images assessing DC viability following treatment with increasing concentrations of OVA. (**b**) Quantitative analysis of DC viability under varying OVA concentrations (n = 4, mean ± s.d.). (**c**) Western blot analysis of MHC expression in DCs treated with OVA.

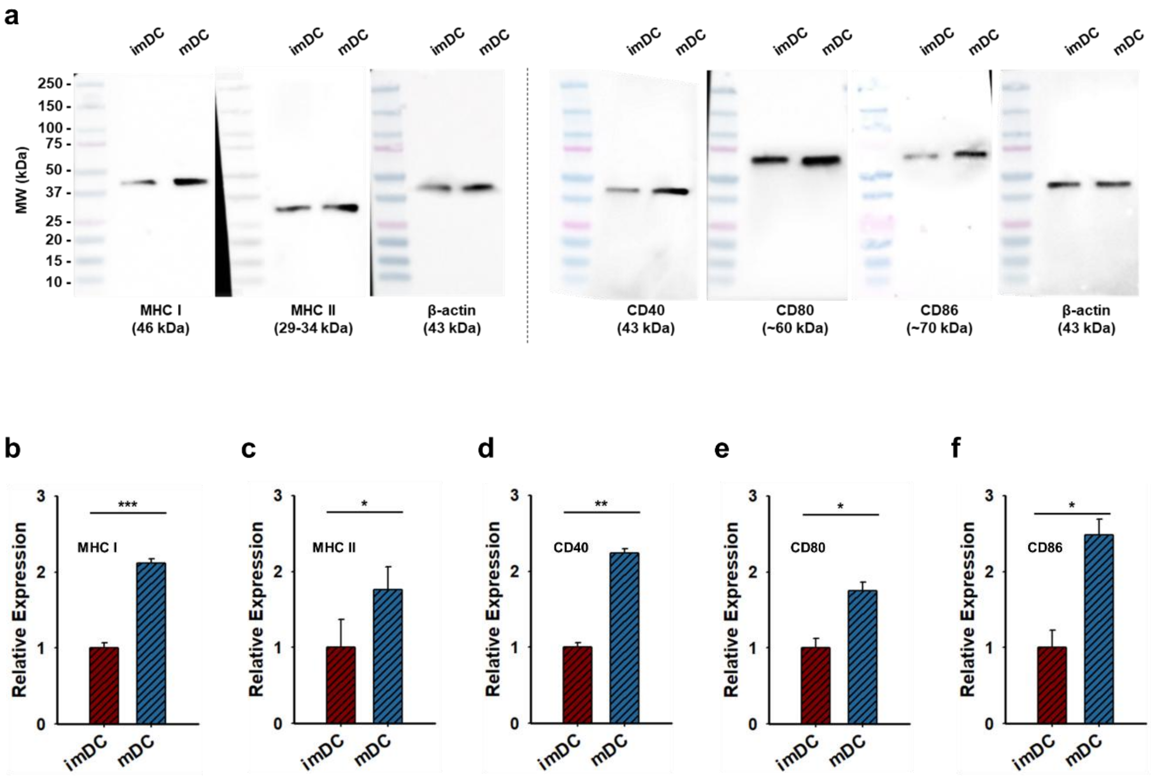


**Supplementary Fig. 2:** Comparison of protein expression between immature dendritic cell (imDC; non-OVA-stimulated) and mature dendritic cell (mDC; OVA-stimulated). (**a**) Full-length Western blot membranes. Relative protein expression levels of (**b**) MHC I, (**c**) MHC II, (**d**) CD40, (**e**) CD80, and (**f**) CD86 (n = 3, mean ± s.d.).

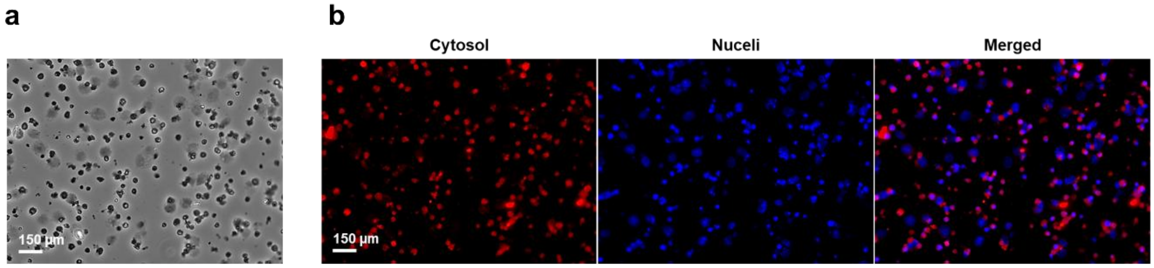


**Supplementary Fig. 3:** Representative images of OT-1 mouse–derived CD8$^{+}$ T cells captured under (**a**) brightfield, and (**b**) fluorescence microscopy. Each channel represents: red for the cytosol and blue for nuclei.

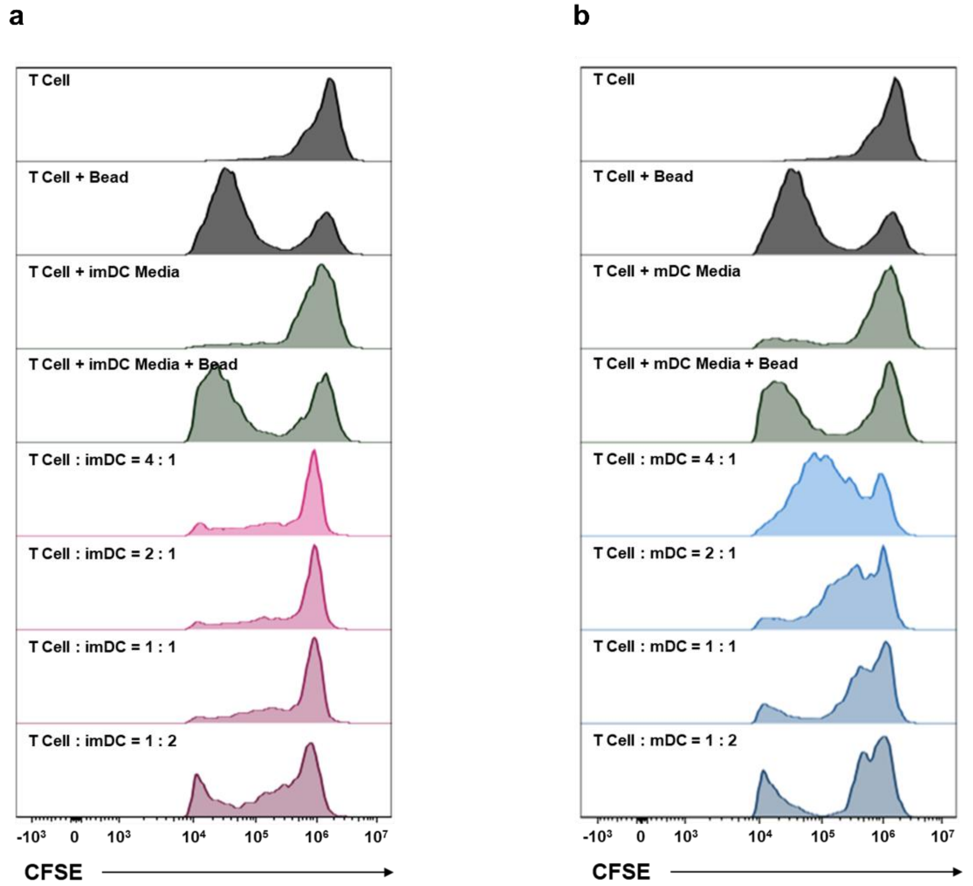


**Supplementary Fig. 4:** T cell proliferation assay. CFSE-labeled T cells were co-cultured with (**a**) immature dendritic cells (imDC) and (**b**) mature dendritic cells (mDC).

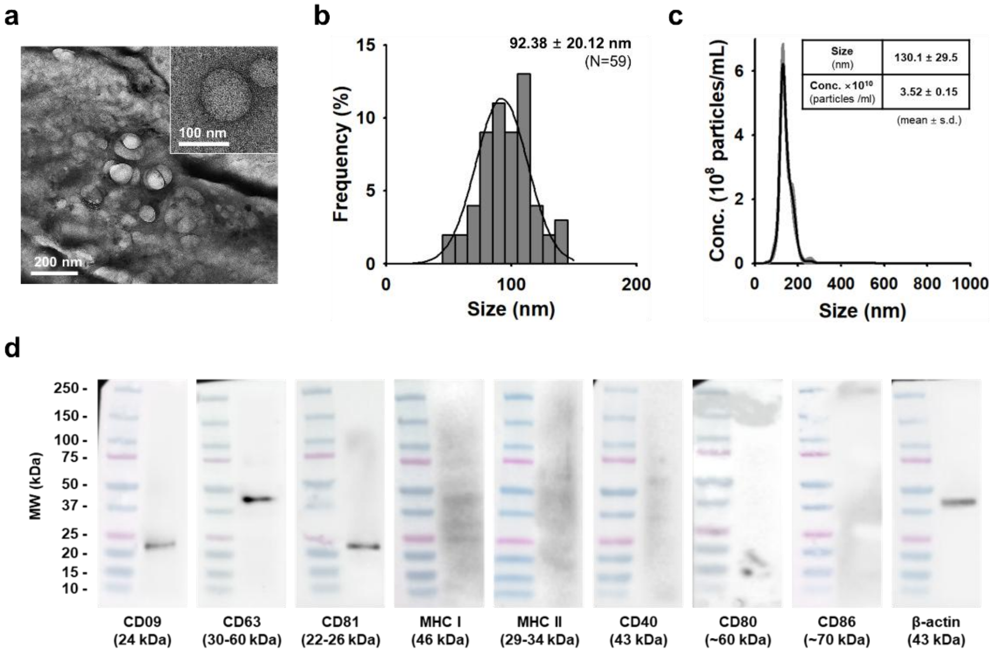


**Supplementary Fig. 5:** Characterization of 3T3-sEVs. (**a**) Representative TEM images showing spherical nanostructures, and (**b**) corresponding size distribution analysis (n=59; mean ± s.d). (**c**) Size distribution and particle concentration profile (n=3; mean ± s.d) measured by NTA. (**d**) Full-length Western blot membranes of sEV markers, MHC molecules, and costimulatory ligands.

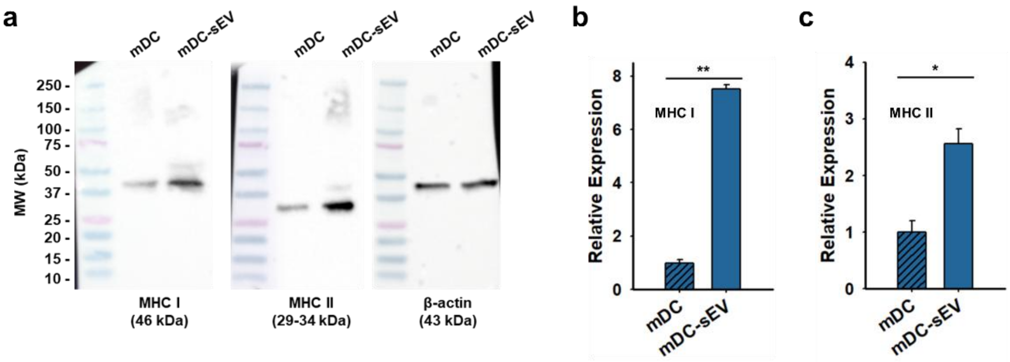


**Supplementary Fig. 6:** Comparison of protein expression between mDC and mDC-sEV. **(a)** Full-length Western blot membranes. Relative protein expression levels of (**b**) MHC I, and **(c)** MHC II (n = 3, mean ± s.d.).

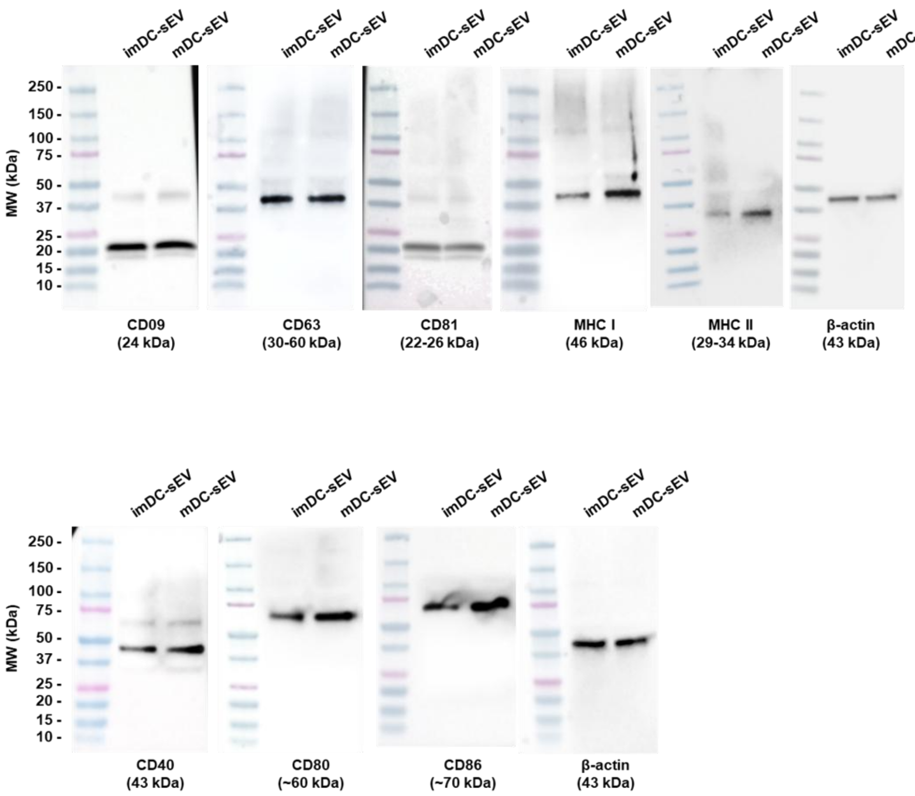


**Supplementary Fig. 7:** Full-length Western blot membranes of sEV markers, MHC, and costimulatory ligands.

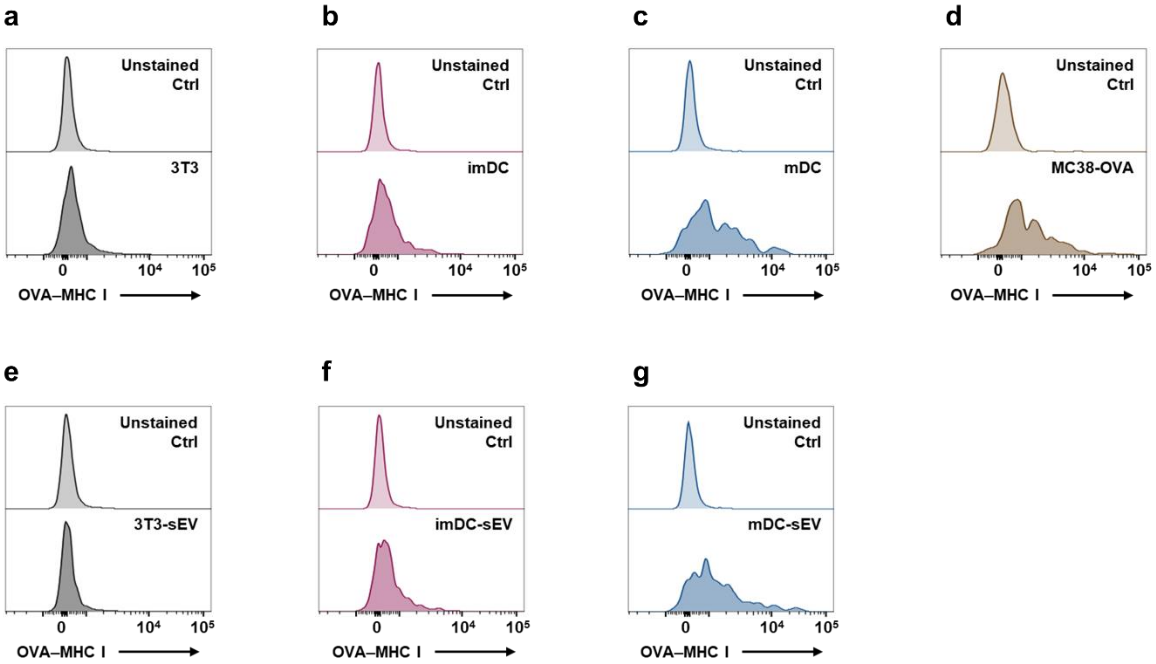


**Supplementary Fig. 8:** Detection of OVA-specific MHC class I (OVA–MHC I; SIINFEKL/H-2K$^{b}$) complexes by flow cytometry. Cell line analysis: (**a**) 3T3 cells, (**b**) imDCs, (**c**) mDCs, and (**d**) MC38-OVA cells. sEV analysis: (**e**) 3T3-derived sEVs, (**f**) imDC-derived sEVs, and (**g**) mDC-derived sEVs.

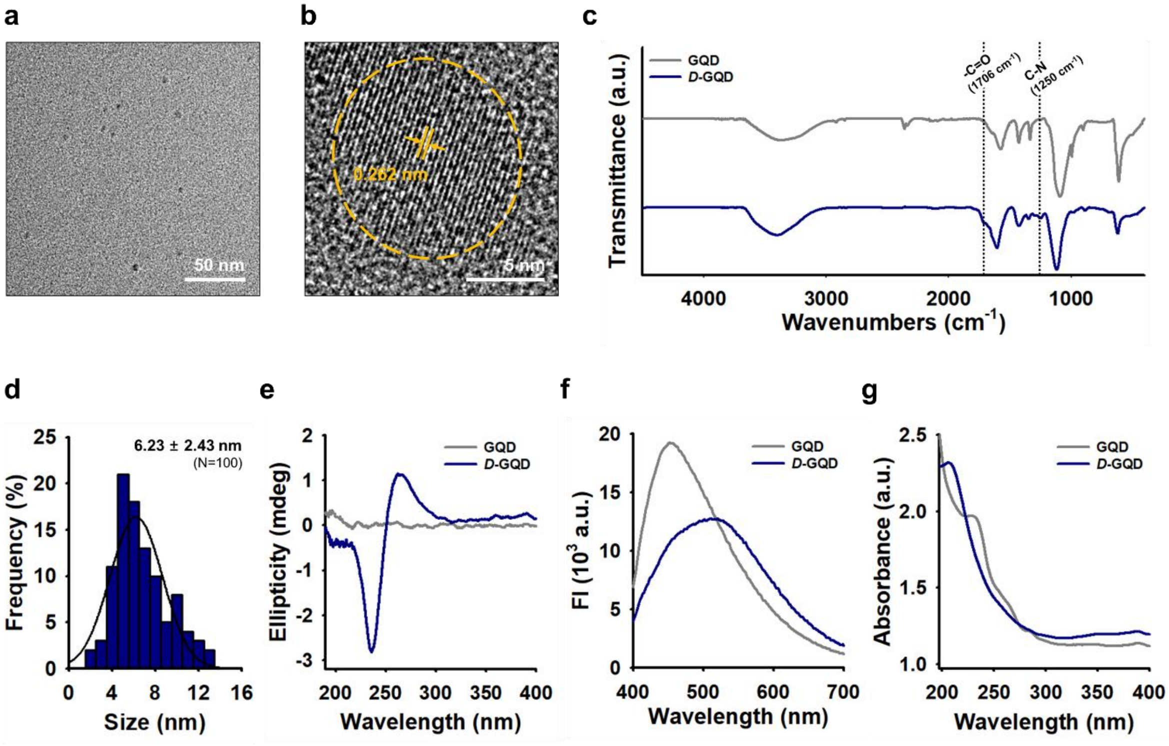


**Supplementary Fig. 9:** Characterization of chiral graphene quantum dots (GQDs). (**a**) TEM image of *D*-GQDs, and (**b**) High-resolution image showing the crystalline lattice structure. (**c**) FTIR spectra. (**d**) size distribution of *D*-GQDs based on TEM image analysis (n = 100, mean ± s.d.). (**e**) Circular dichroism spectra. (**f**) Fluorescence emission spectra excited at 360 nm. (**g**) UV–vis absorption spectra.

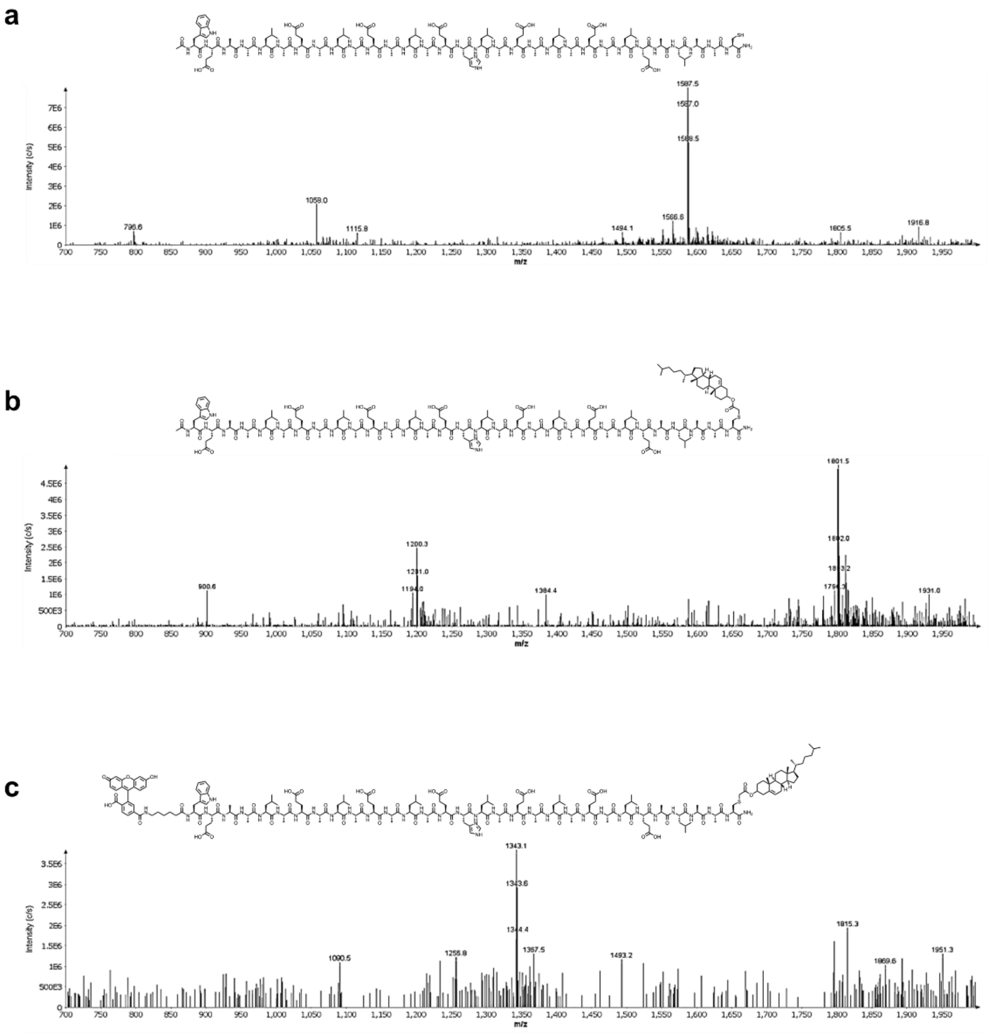


**Supplementary Fig. 10:** ESI-MS verification of (**a**) GALA peptide, (**b**) modified GALA-chol peptide, and **c**, modified FAM-GALA-chol peptide.

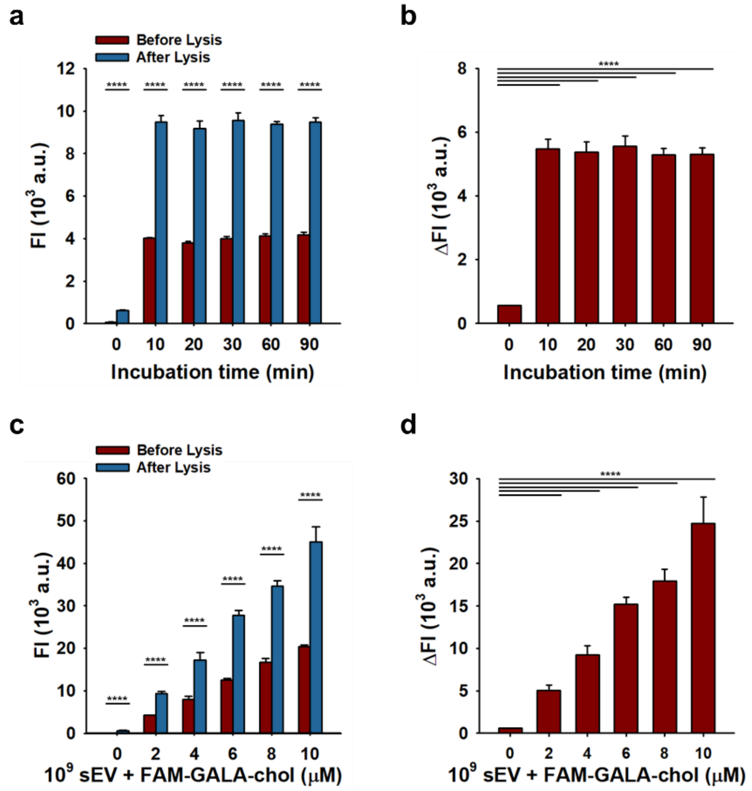


**Supplementary Fig. 11:** Characterization of GALA-functionalized mDC-sEVs. Fluorescence recovery of FAM labeled on GALA-chol before and after sEV membrane lysis. (**a** and **b**) optimization of co-incubation time. (**c** and **d**) concentration-dependent incorporation. (n = 4, mean ± s.d.). Ex/Em = 490/520 nm.

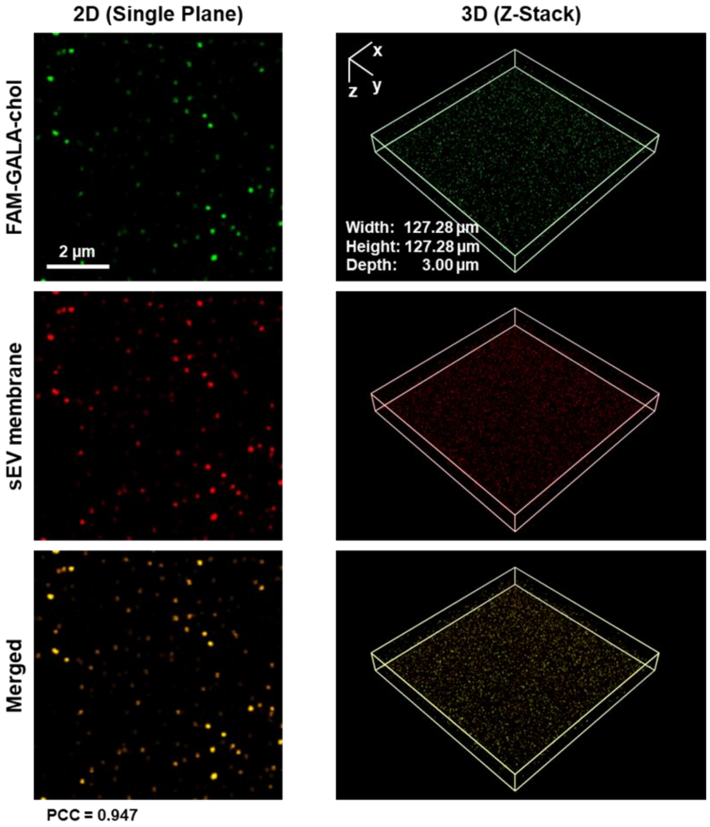


**Supplementary Fig. 12:** Characterization of GALA functionalized mDC-sEVs under CLSM. Each CLSM channel represents: green for FAM-GALA-chol and red for sEV membrane.

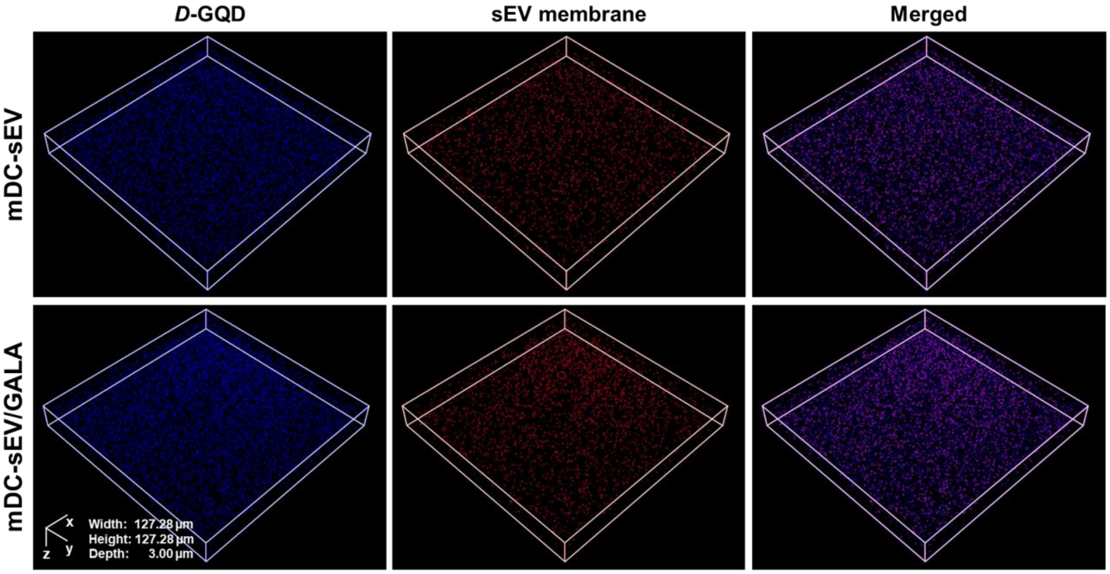


**Supplementary Fig. 13:** Z-stack characterization of mDC-sEVs loaded with *D*-GQDs and surface-functionalized with the GALA peptide under CLSM. Each CLSM channel represents: blue for *D*-GQDs and red for sEV membrane.

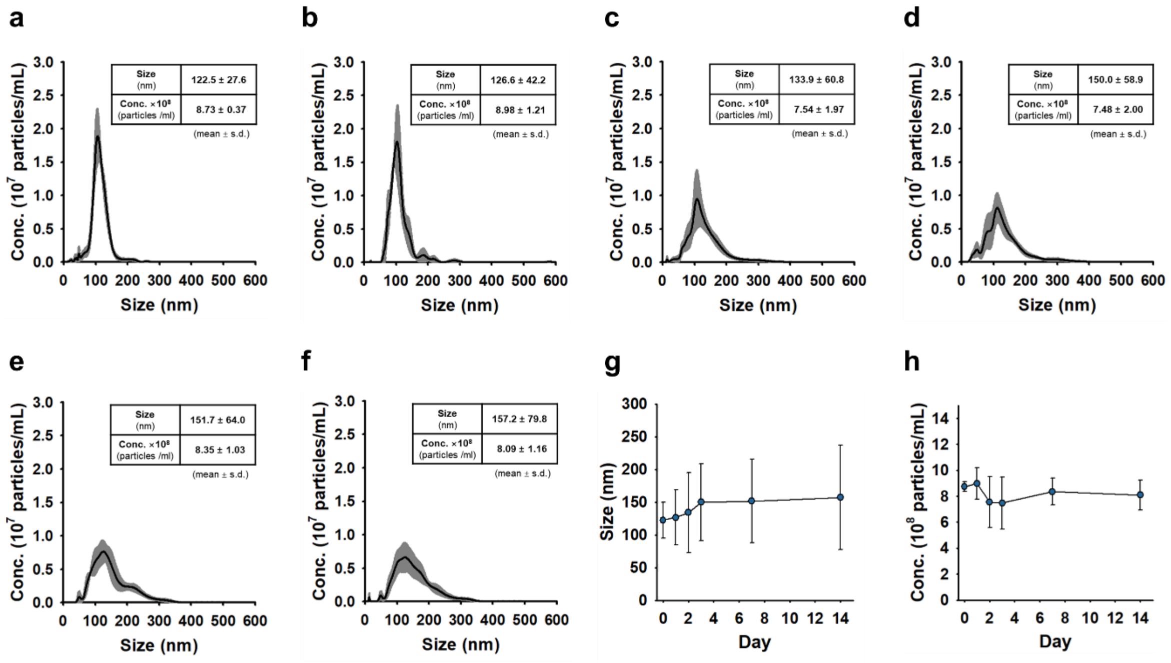


**Supplementary Fig. 14:** Time-dependent structural integrity assessment of *D*-GQD–loaded, GALA-functionalized mDC-sEVs by NTA measurements at (**a**) 0; (**b**) 1; (**c**) 2; (**d**) 3; (**e**) 7; and (**f**) 14 days. (**g**) Comparison of time-dependent hydrodynamic size changes. (**h**) Comparison of time-dependent particle concentration changes.

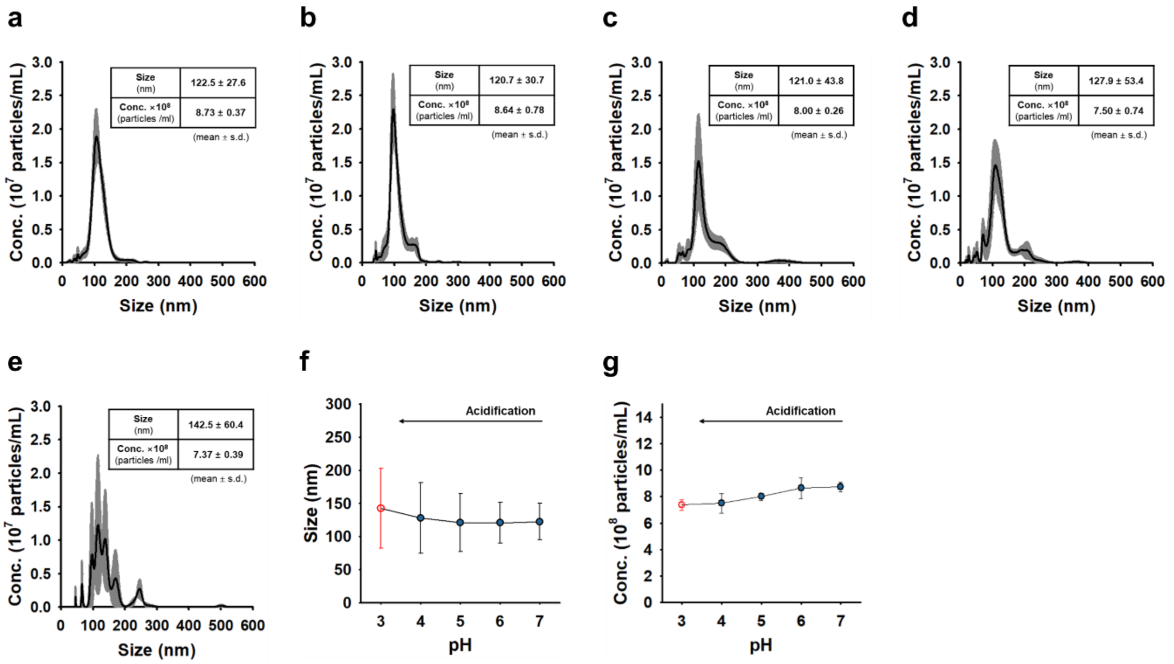


**Supplementary Fig. 15:** pH-dependent structural integrity assessment of *D*-GQD–loaded, GALA-functionalized mDC-sEVs by NTA measurements at pH (**a**) 7; (**b**) 6; (**c**) 5; (**d**) 4; and (**e**) 3. (**f**) Comparison of pH-dependent hydrodynamic size changes. (**g**) Comparison of pH-dependent particle concentration changes.

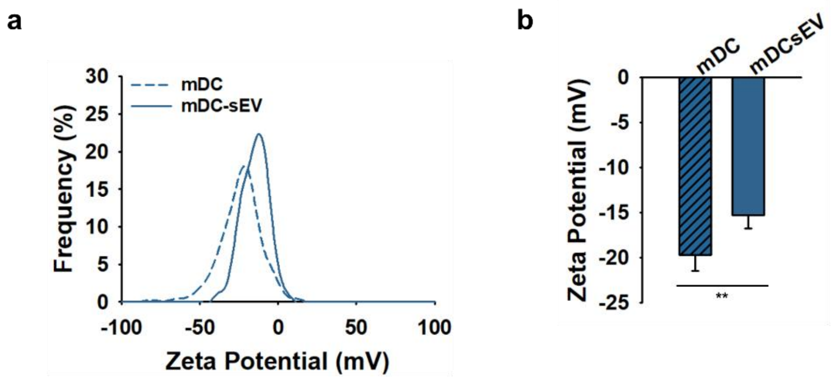


**Supplementary Fig. 16:** Zetasizer analysis of mDCs and mDC-sEVs at pH 7.0. (**a**) Zeta potential distributions. (**b**) Comparison of average surface charge (n = 4; mean ± s.d.).

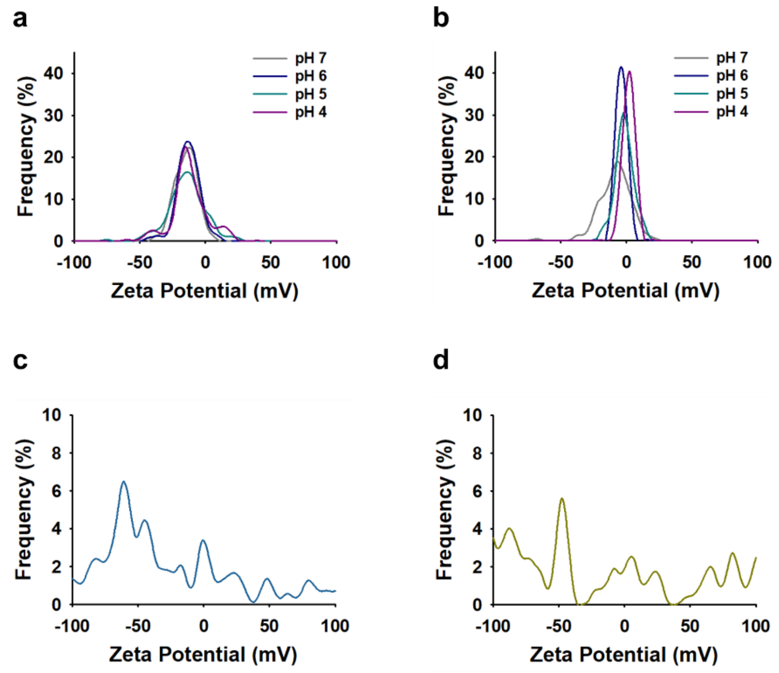


**Supplementary Fig. 17:** Zetasizer analysis of GALA-functionalized mDC-sEVs under varying pH conditions. Zeta potential distributions of (**a**) non-functionalized $10^9$ sEVs and **b**, GALA–chol-functionalized $10^9$ sEVs. Zeta potential distributions at pH 3 for **c**, non-functionalized sEVs and (**d**) GALA-functionalized sEVs.

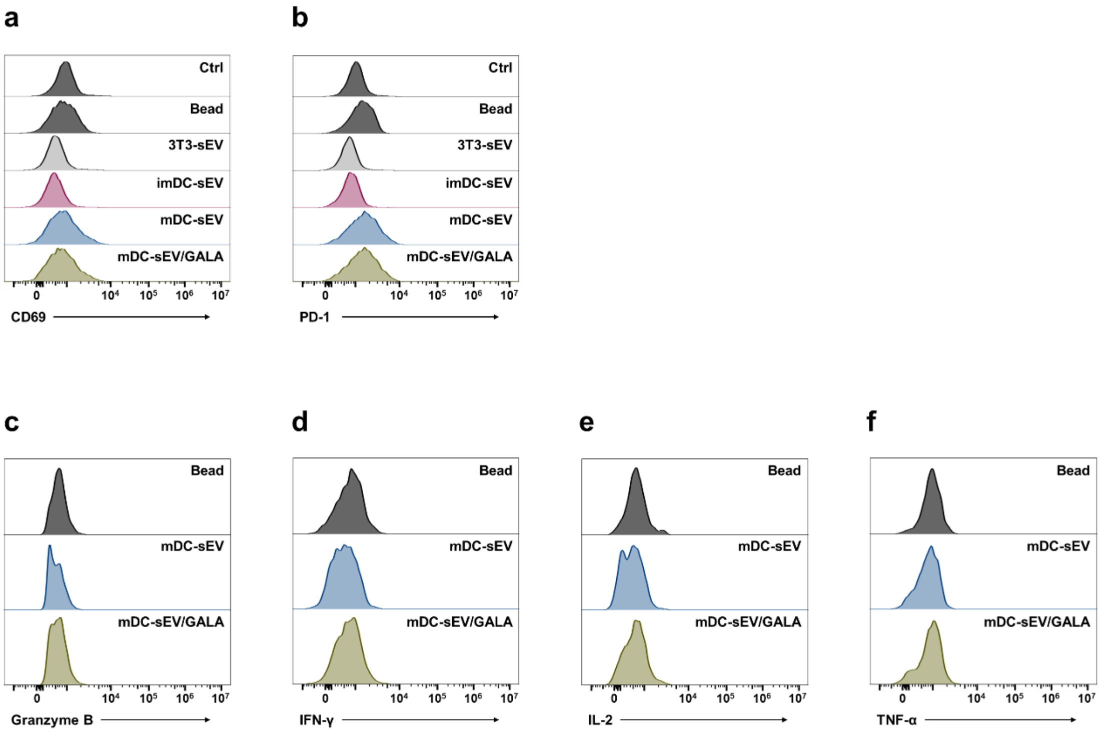


**Supplementary Fig. 18:** Flow cytometry unstained controls. Assessment of T-cell activation by the proportions of (**a**) $CD8^+/CD69^+$, and (**b**) $CD8^+/PD\text{-}1^+$ cells. Assessment of activated T-cell–mediated intracellular cytokine production by the proportions of (**c**) $CD8^+$/Granzyme $B^+$, (**d**) $CD8^+/IFN\text{-}\gamma^+$, (**e**) $CD8^+/IL\text{-}2^+$, and (**f**) $CD8^+/TNF\text{-}\alpha^+$ cells.

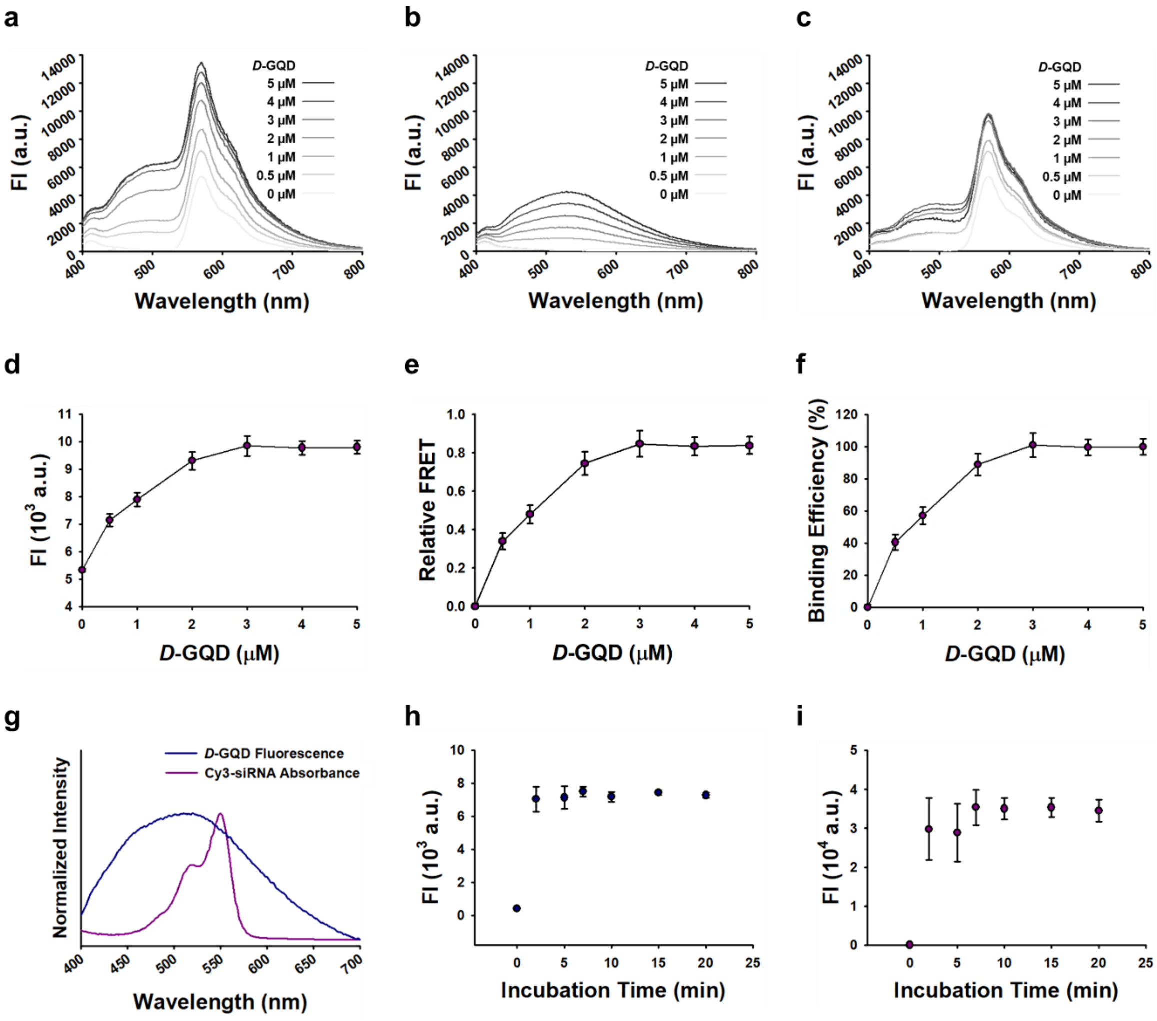


**Supplementary Fig. 19:** Characterization of siRNA/*D*-GQD complexes. (**a**) Fluorescence spectra of Cy3-labeled siRNA (5 μM) incubated with increasing concentrations of *D*-GQDs (0–5 μM) under excitation at 360 nm (*D*-GQD excitation wavelength). (**b**) Fluorescence spectra of *D*-GQDs at varying concentrations (0–5 μM). (**c**) *D*-GQD background-subtracted fluorescence spectra of siRNA/*D*-GQD complexes after deduction of *D*-GQD intrinsic fluorescence. (**d**) Fluorescence intensity of siRNA/*D*-GQD complexes at 570 nm (Cy3 emission wavelength) upon excitation at 360 nm. (**e**) Relative FRET efficiency, and (**f**) corresponding siRNA binding efficiency to *D*-GQDs. (**g**) Spectral overlap between *D*-GQD excitation and Cy3-siRNA absorbance used for FRET analysis. Optimization of incubation time for (**h**) siRNA/*D*-GQD complexation, and (**i**) loading of siRNA/*D*-GQD complexes into sEVs. (n= 3, mean ± s.d.).

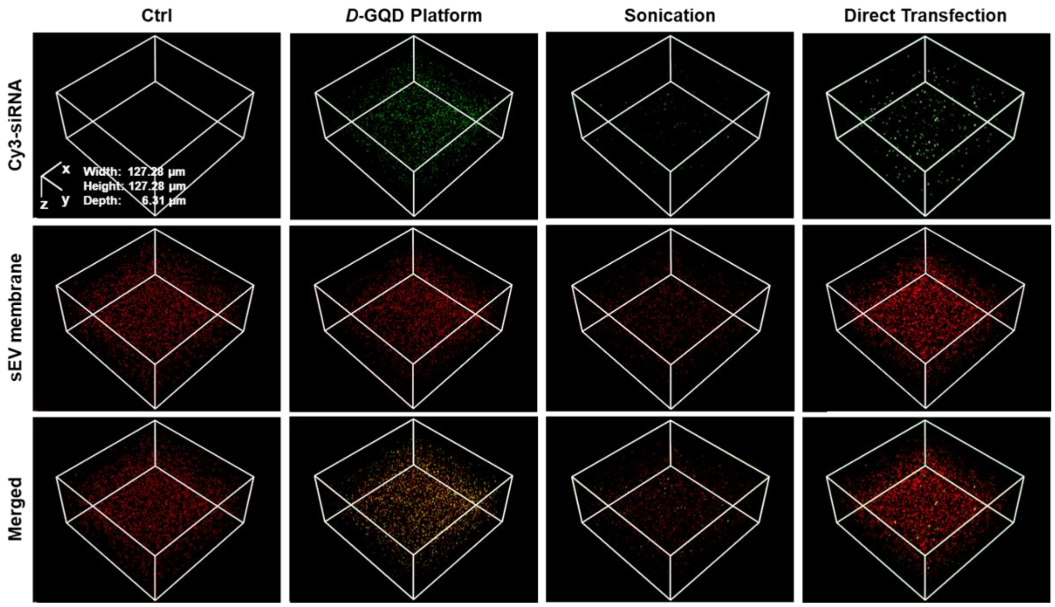


**Supplementary Fig. 20:** Z-stack characterization of mDC-sEVs loaded with siRNA under CLSM. siRNA was loaded via *D*-GQD chiral-assisted loading, sonication, or direct transfection using Lipofectamine. Each CLSM channel represents: green for Cy3-siRNA and red for sEV membrane.

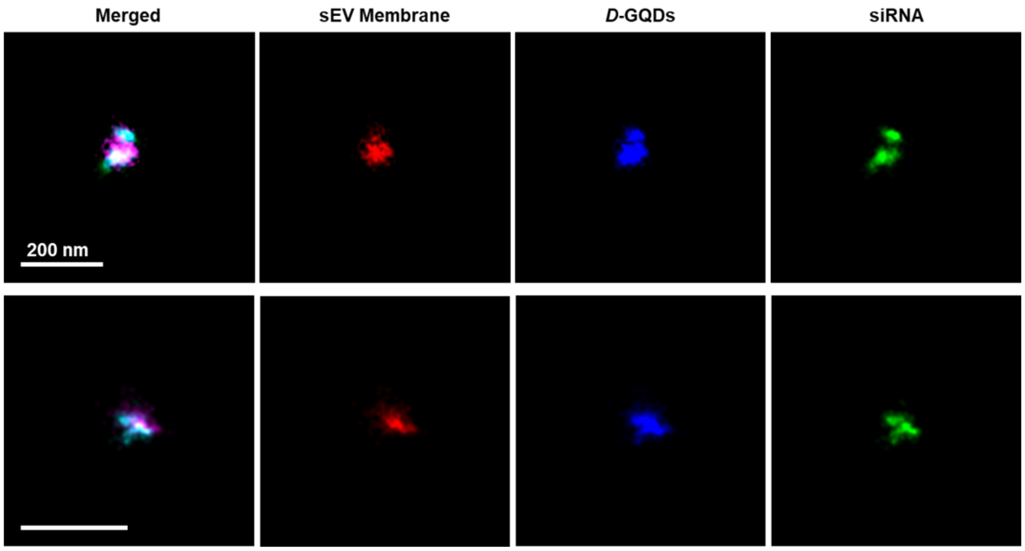


**Supplementary Fig. 21:** Super-resolution images of individual mDC-sEVs obtained by single-molecule localization microscopy. Each channel represents: red for sEV membrane, blue for *D*-GQD, and green for Cy3-siRNA.

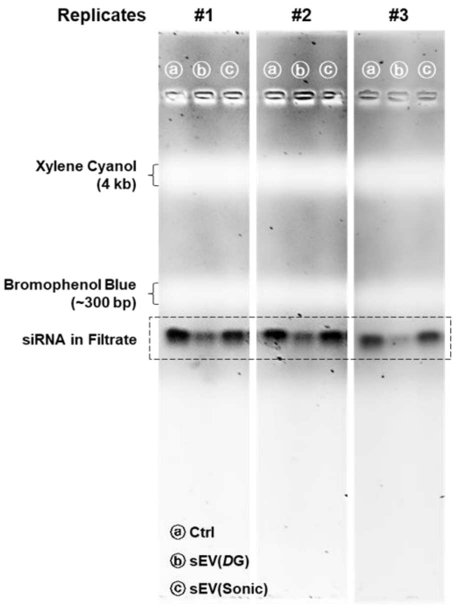


**Supplementary Fig. 22:** Full-length agarose gel after electrophoresis for quantification of unincorporated siRNA after loading via *D*-GQD (*D*G) and Sonic approaches.

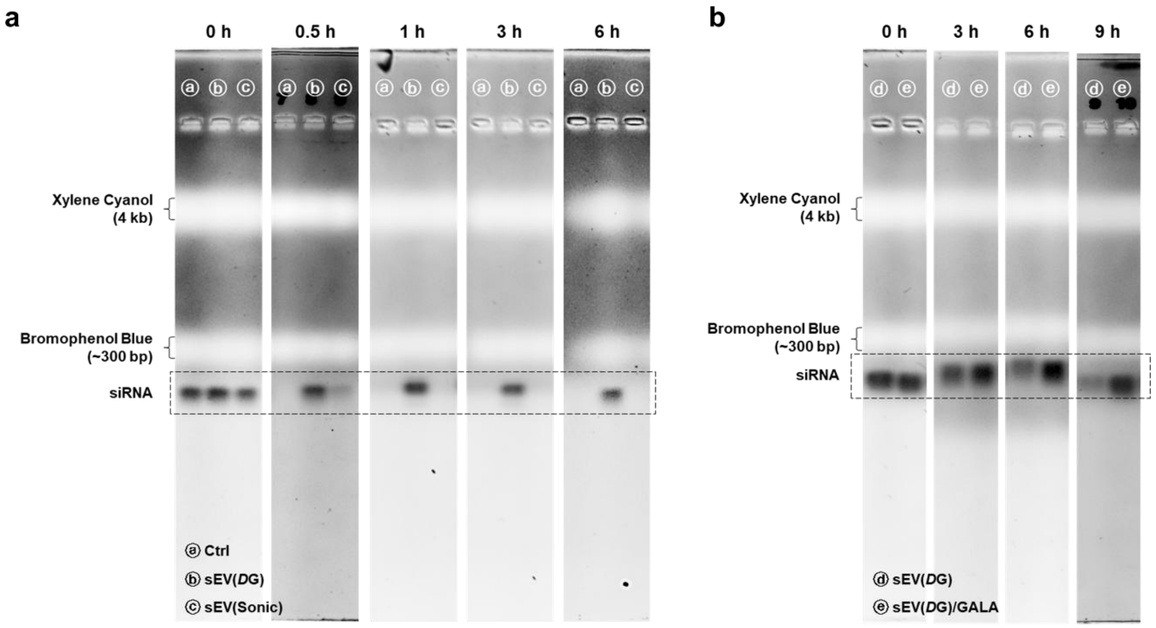


**Supplementary Fig. 23:** Full-length agarose gel after electrophoresis. (**a**) Time-dependent siRNA integrity comparison for different siRNA loading approaches into mDC-sEVs. *D*G: *D*-GQD chiral-assisted loading; Sonic: sonication loading. (**b**) Time-dependent siRNA integrity comparison before and after GALA functionalization of the mDC-sEV surface.

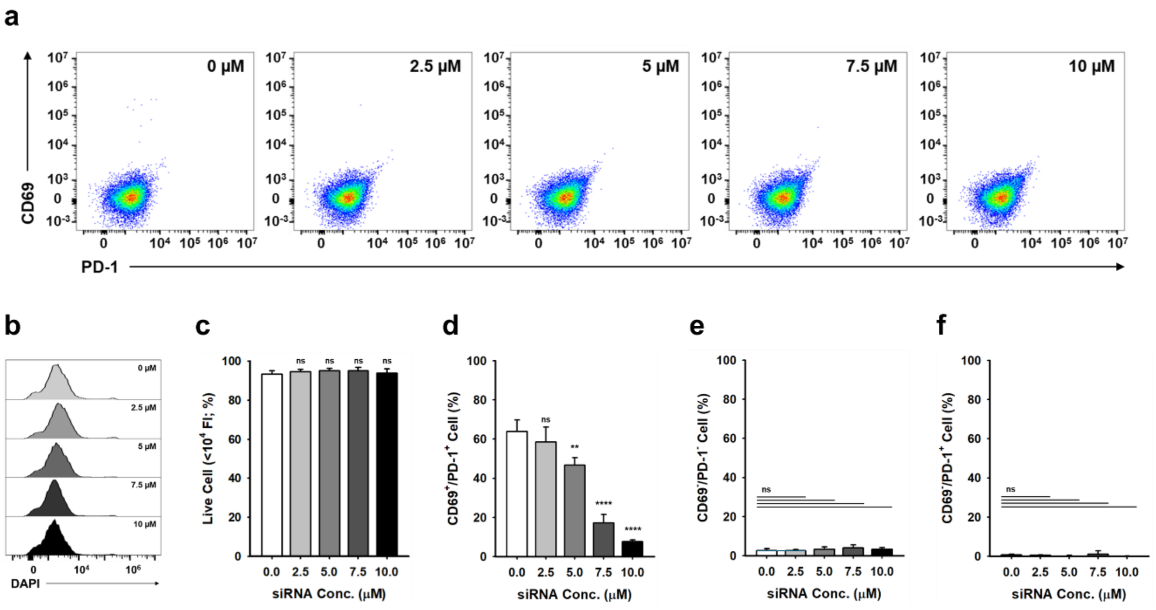


**Supplementary Fig. 24:** Dose-dependent PD-1 silencing efficacy test of mDC-sEV(siR) treated CD8$^+$ T cells. (**a**) Flow cytometry unstained controls. (**b** and **c**) siRNA concentration-dependent cell viability via quantification of DAPI-negative cells. Assessment of T-cell activation by the proportions of (**d**) CD69$^+$/PD-1$^+$, (**e**) CD69$^-$/PD-1$^-$, and (**f**) CD69$^-$/PD-1$^+$ cells. 0 µM groups: treatment of mDC-sEV without siRNA loading.

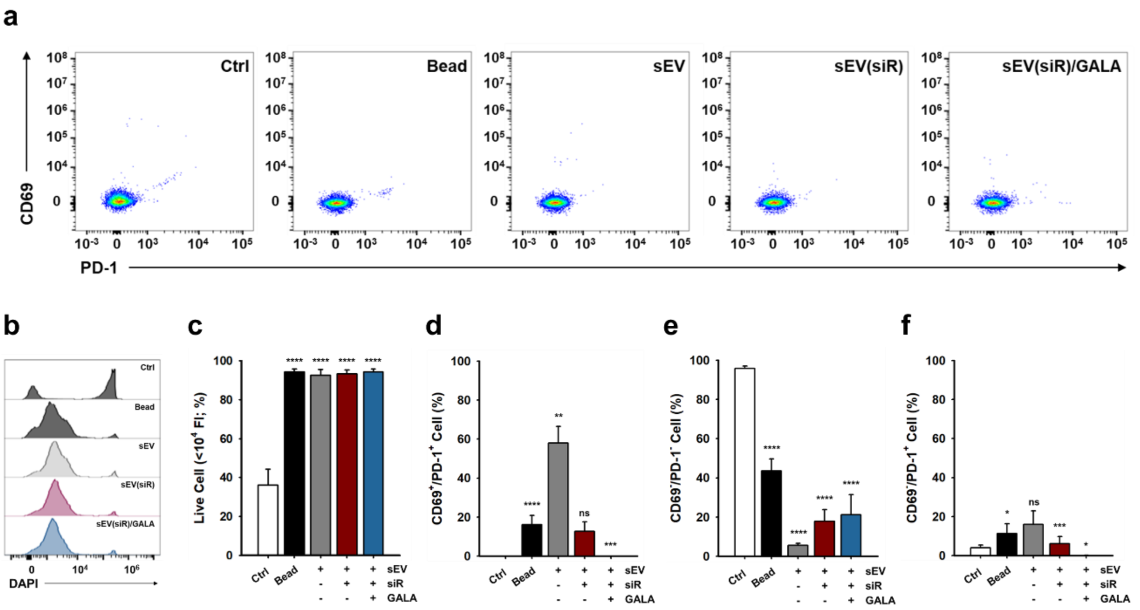


**Supplementary Fig. 25:** Comparative PD-1 silencing efficacy test of mDC-sEV(siR) treated CD8$^{+}$ T cells. (**a**) Flow cytometry unstained controls. (**b** and **c**) treatment type-dependent cell viability via quantification of DAPI-negative cells. Assessment of T-cell activation by the proportions of (**d**) CD69$^{+}$/PD-1$^{+}$, (**e**) CD69$^{-}$/PD-1$^{-}$, and (**f**) CD69$^{-}$/PD-1$^{+}$ cells. Ctrl groups: non-treatment (PBS).

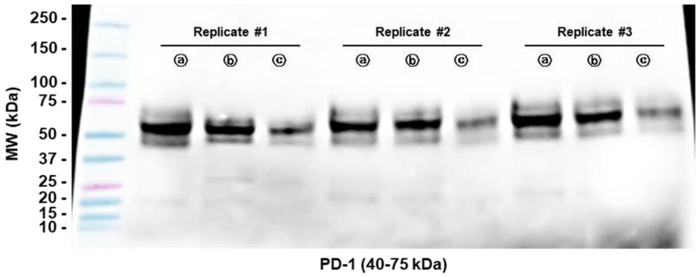


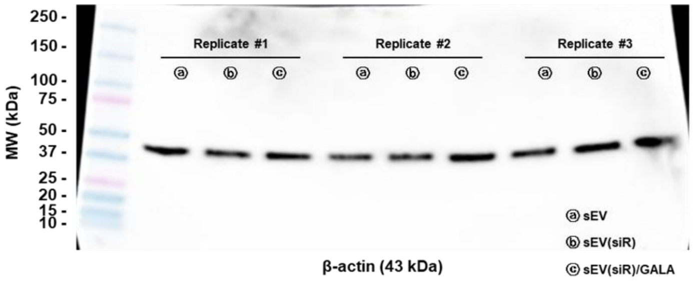


**Supplementary Fig. 26:** Full-length Western blot membranes of PD-1 expression in $CD8^+$ T cells following siRNA transfection via mDC-sEVs.

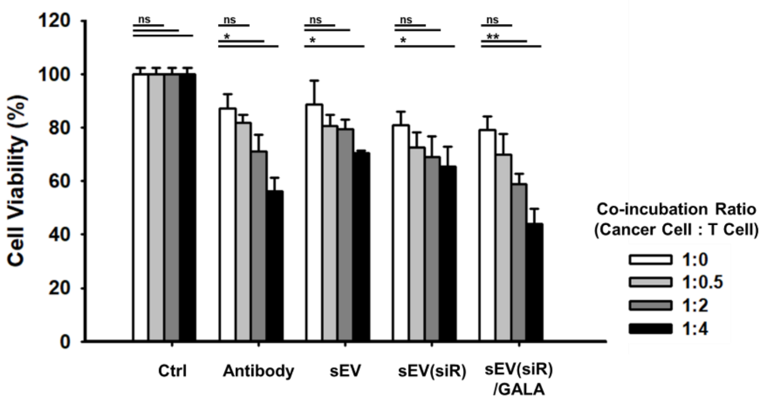


**Supplementary Fig. 27:** Quantification of co-incubation ratio–dependent *in vitro* therapeutic effects. Cancer cell viability test following co-culture with activated T cells treated with Ctrl, Antibody, sEV, sEV(siR), or sEV(siR)/GALA. Cancer cell: MC38-OVA cell line; sEV type: mDC-sEV. (n = 3, mean ± s.d.).

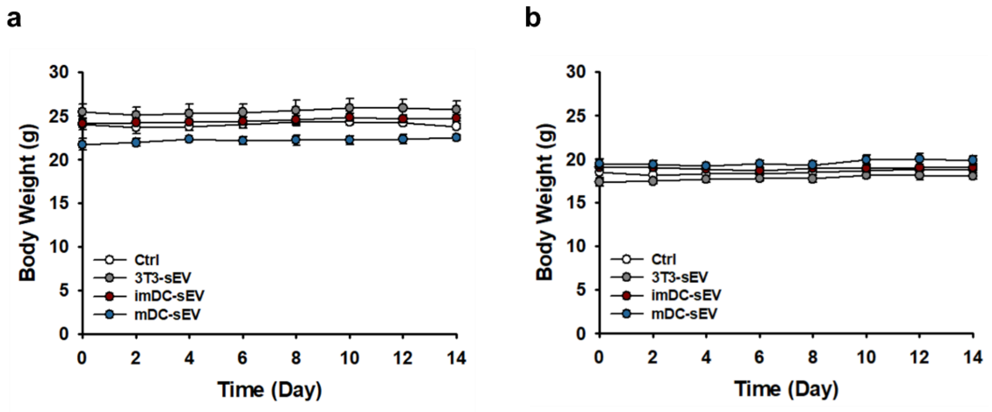


**Supplementary Fig. 28:** Safety evaluation of sEVs by body weight monitoring of mice following tail-vein i.v. injection. (**a**) Male and (**b**) female mice. (n = 3, mean ± s.d.).

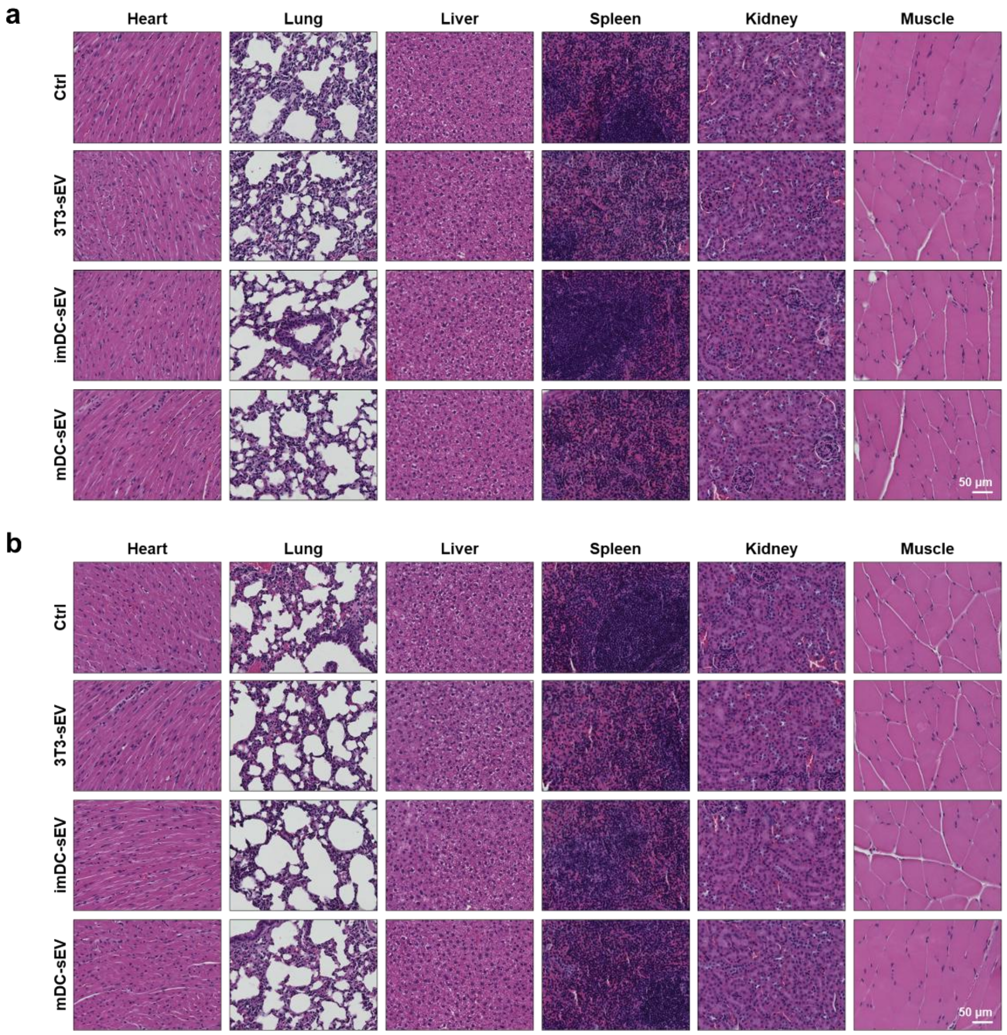


**Supplementary Fig. 29:** Safety evaluation of sEVs by H&E-stained histological analysis of major organs collected two weeks post-injection following tail-vein i.v. injection. (**a**) Male and (**b**) female mice.

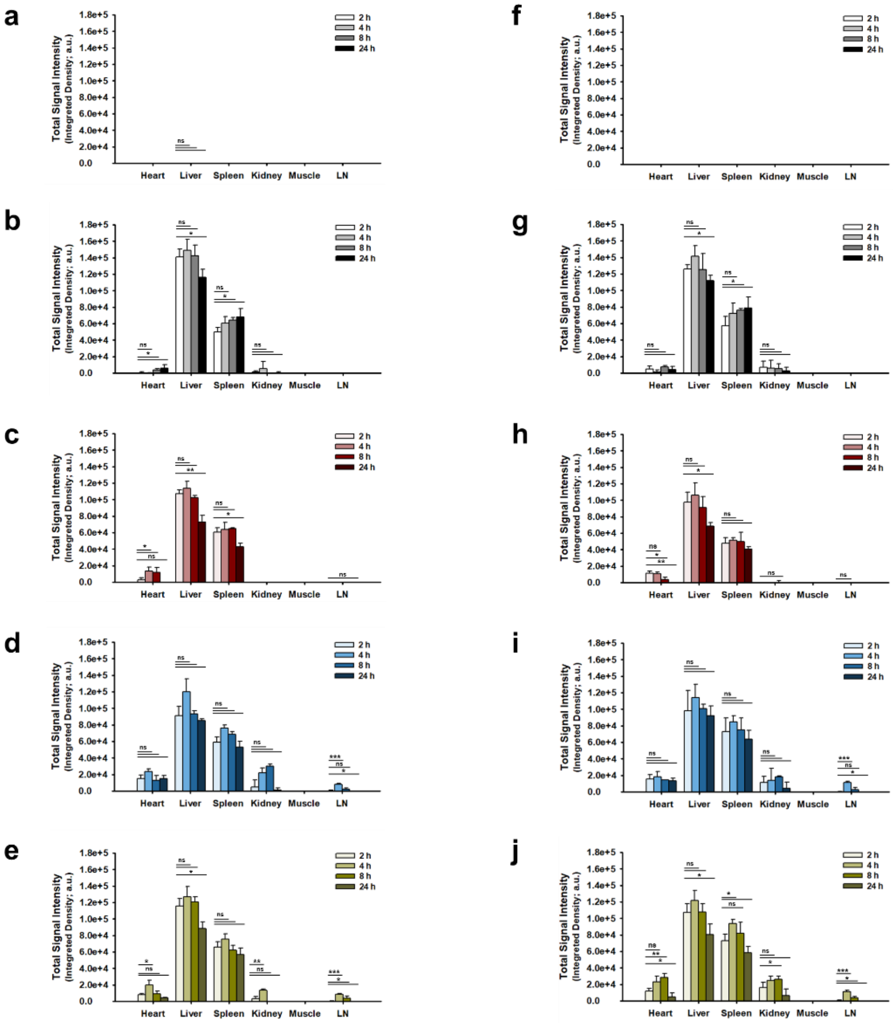


**Supplementary Fig. 30:** Biodistribution quantification of *ex vivo* fluorescence images of major organs resected from mice administered with male mice (**a**) PBS, (**b**) 3T3-sEVs, (**c**) imDC-sEVs, (**d**) mDC-sEVs, and (**e**) mDC-sEVs/GALA and female mice (**f**) PBS, (**g**) 3T3-sEVs, (**h**) imDC-sEVs, (**i**) mDC-sEVs, and (**j**) mDC-sEVs/GALA (n = 3, mean ± s.d.). LN: Lymph node.

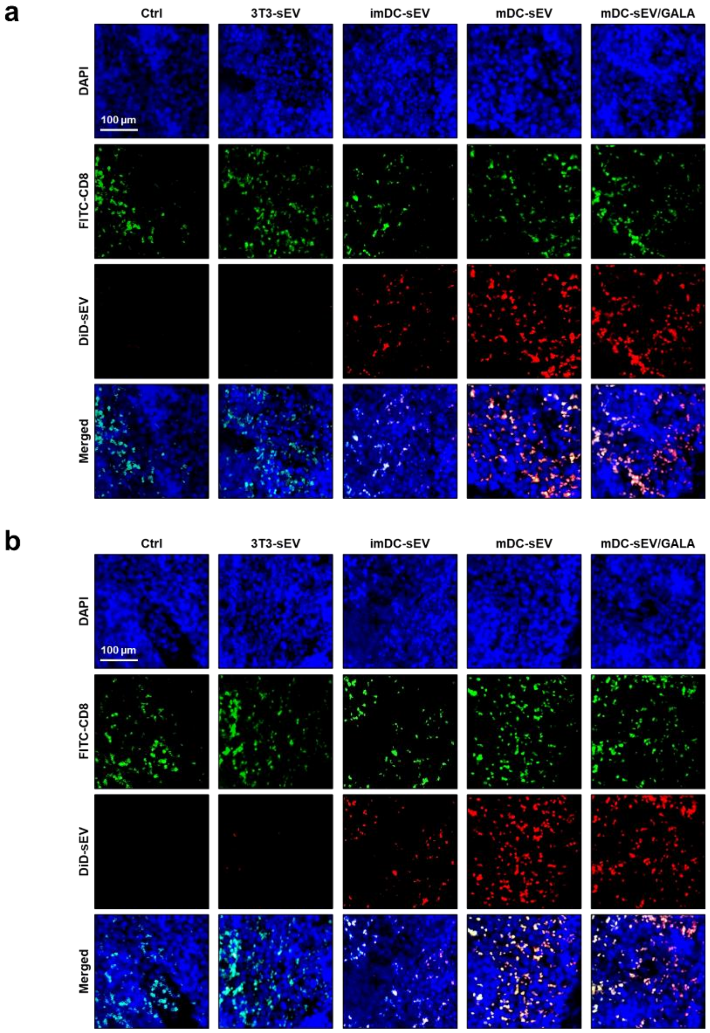


**Supplementary Fig. 31:** Immunofluorescence images of lymph nodes in (**a**) male mice and (**b**) female mice 3 h after tail-vein i.v. injection of DiD-labeled-sEVs. Each channel represents: blue, nuclei; green, $CD8^+$ T cells; red, sEVs.

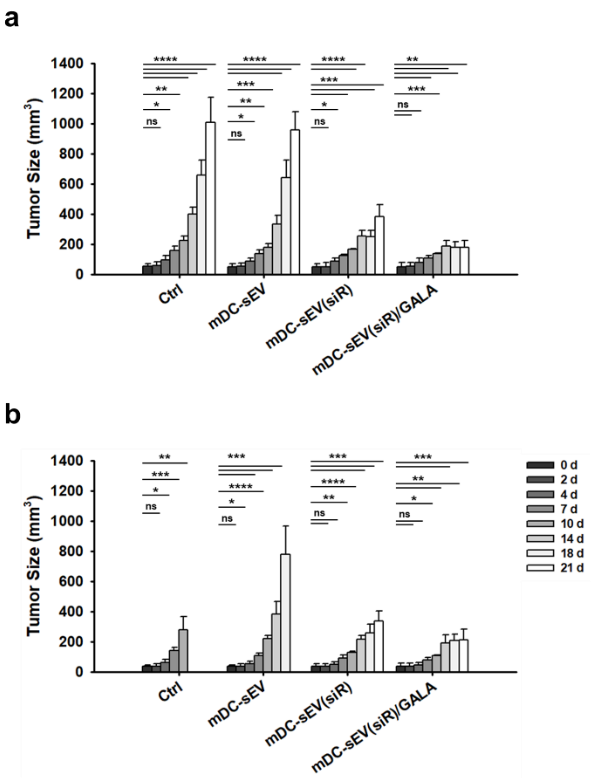


**Supplementary Fig. 32:** Quantification of time-dependent *in vivo* therapeutic effects within groups with statistical analysis for (**a**) male mice and (**b**) female mice (n = 4, mean ± s.d.).

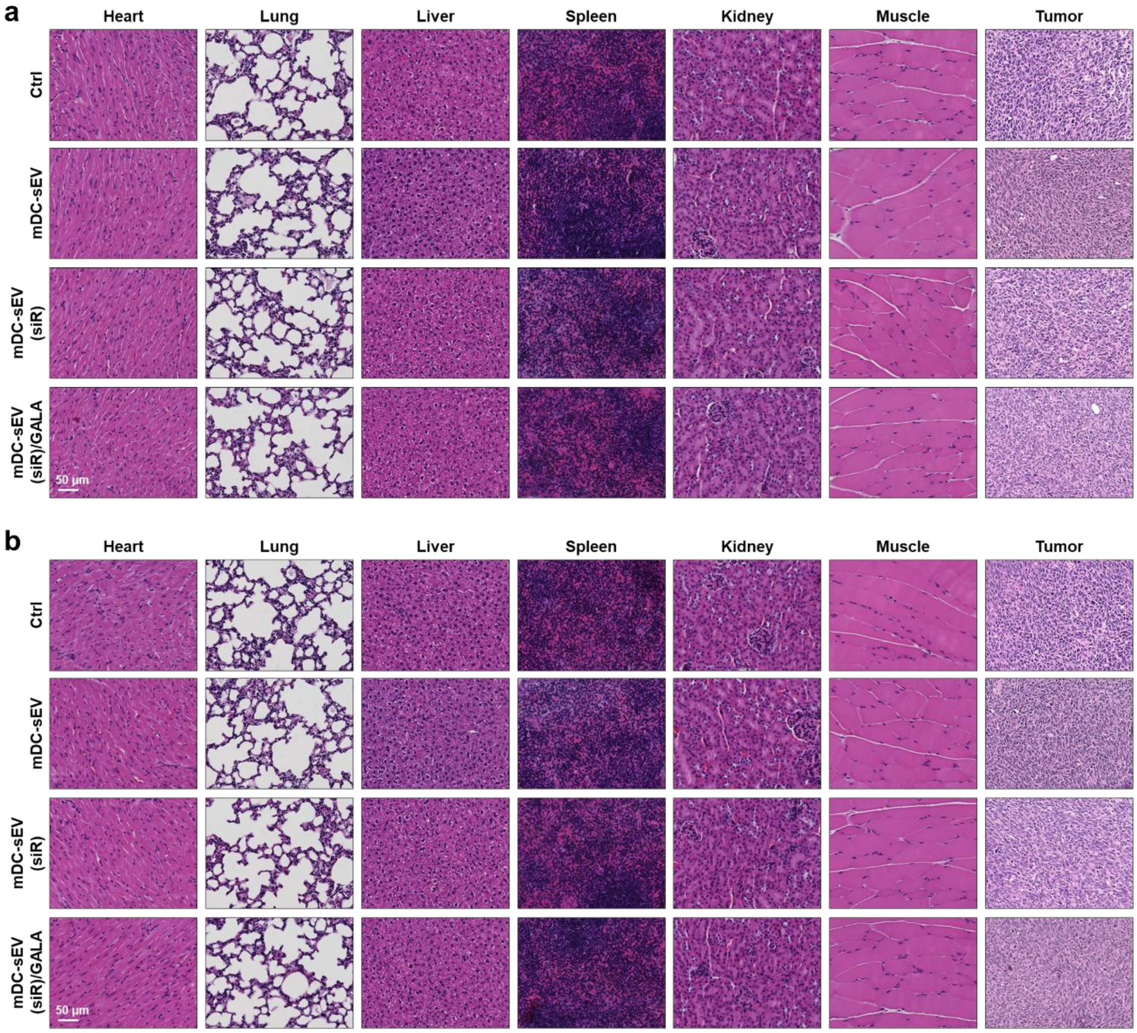


**Supplementary Fig. 33:** H&E-stained histological analysis of major organs and tumor tissues collected at the time of euthanasia for each treatment group following tail-vein intravenous injection. (**a**) male mice and (**b**) female mice.

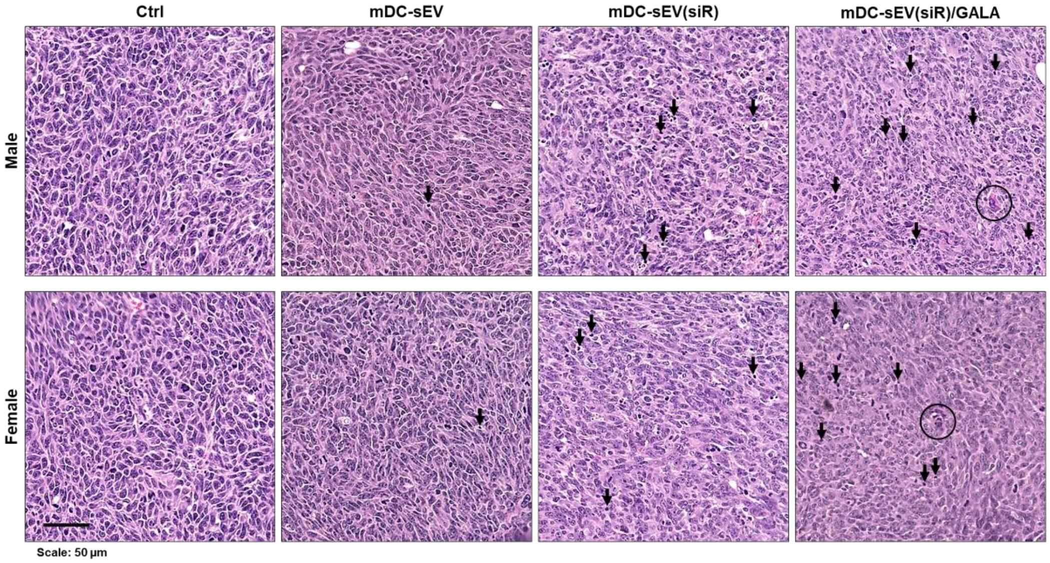


**Supplementary Fig. 34:** H&E-stained histological analysis of magnified tumor sections showing tissue damage. Arrow: tumor-infiltrating lymphocytes (TILs); Circle: TIL-attacked cancer cells.

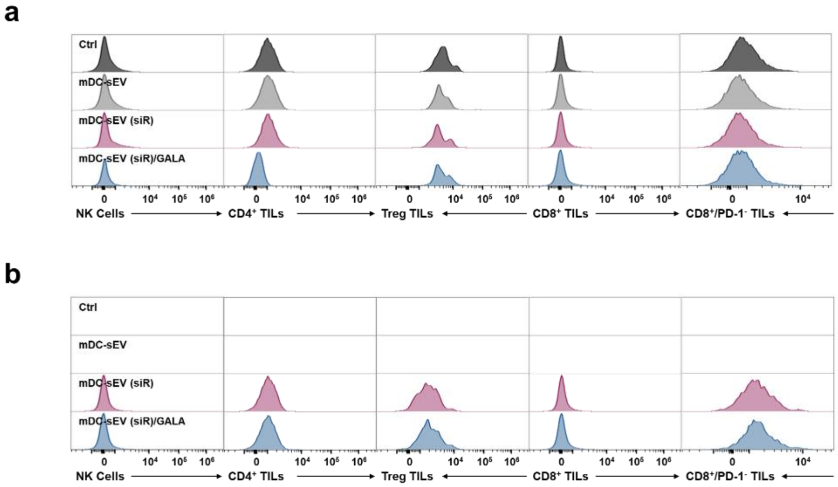


**Supplementary Fig. 35:** TIME reprogramming test after mDC-sEV treatment. Flow cytometry unstained controls of (**a**) male mice and (**b**) female mice.

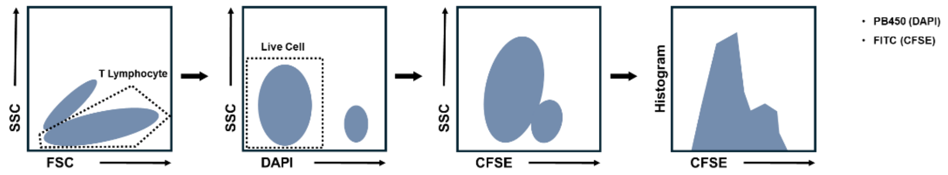


**Supplementary Fig. 36:** Workflow of flow cytometry analysis for the T-cell proliferation assay. Percentage of the population was gated and quantified based on fluorescence intensity <1 × $10^4$.

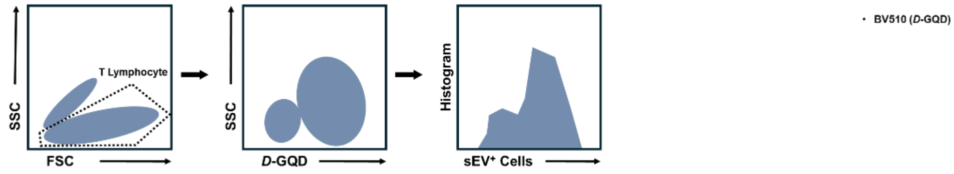


**Supplementary Fig. 37:** Workflow of flow cytometry analysis for sEV uptake into T cells. Percentage of the population was gated and quantified based on fluorescence intensity >5 × $10^3$.

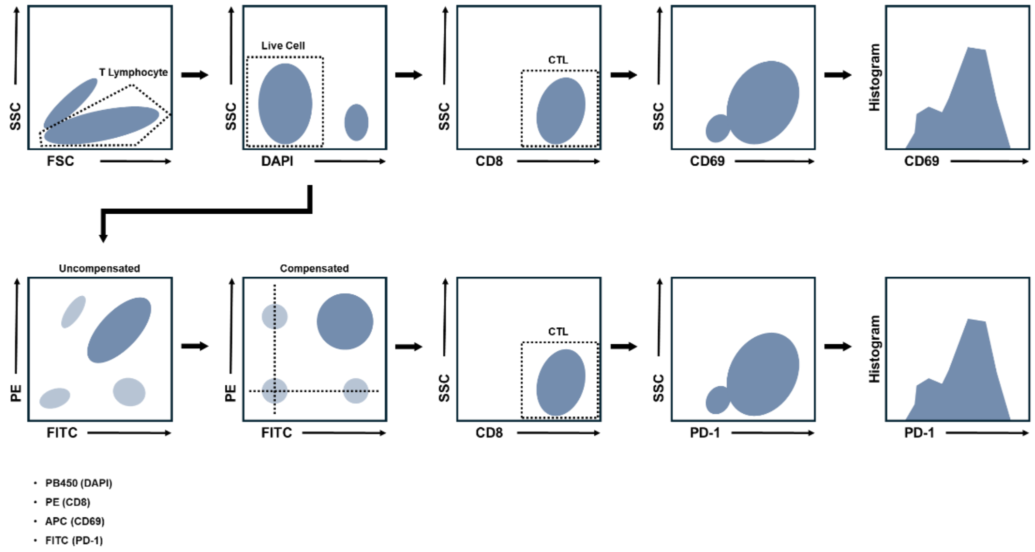


**Supplementary Fig. 38:** Workflow of flow cytometry analysis for T cell activation. Percentage of the population was gated and quantified based on fluorescence intensity >1 × $10^4$.

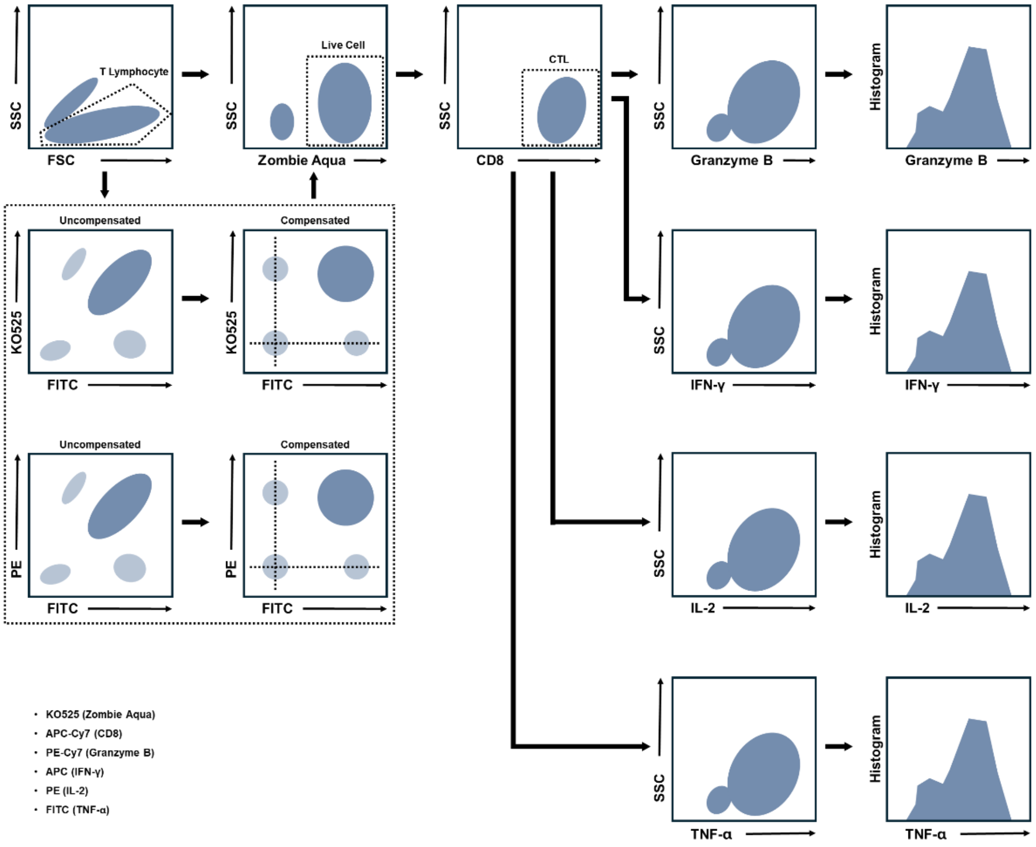


**Supplementary Fig. 39:** Workflow of flow cytometry analysis for activated T-cell–mediated intracellular cytokine production. Percentage of the population was gated and quantified based on fluorescence intensity >5 × $10^3$.

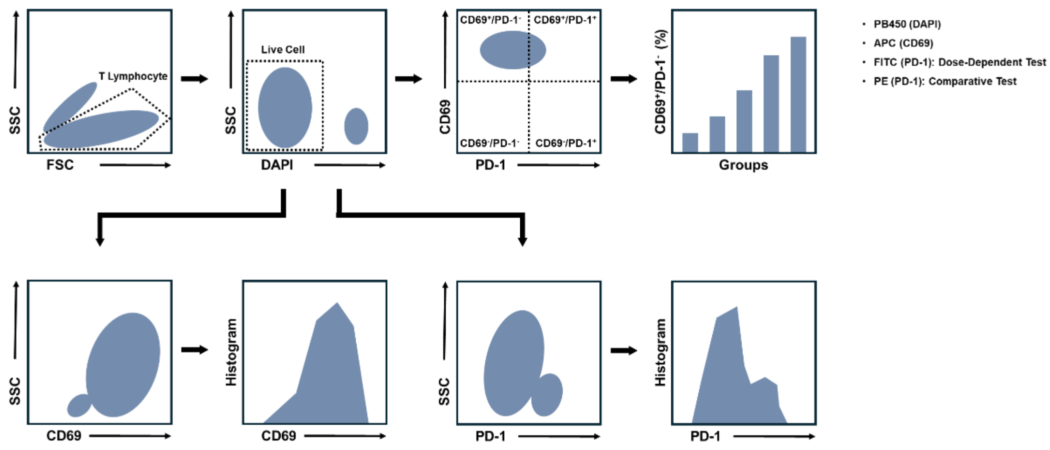


**Supplementary Fig. 40:** Workflow of flow cytometry analysis for PD-1 silencing efficacy test of mDC-sEV(siR) treated $CD8^+$ T cells. Percentage of the population was gated and quantified based on fluorescence intensity $>1 \times 10^4$ for CD69 and $<1 \times 10^4$ for PD-1 in the dose-dependent test; fluorescence intensity $>1 \times 10^5$ for CD69 and $<2 \times 10^3$ for PD-1 in the time-dependent test.

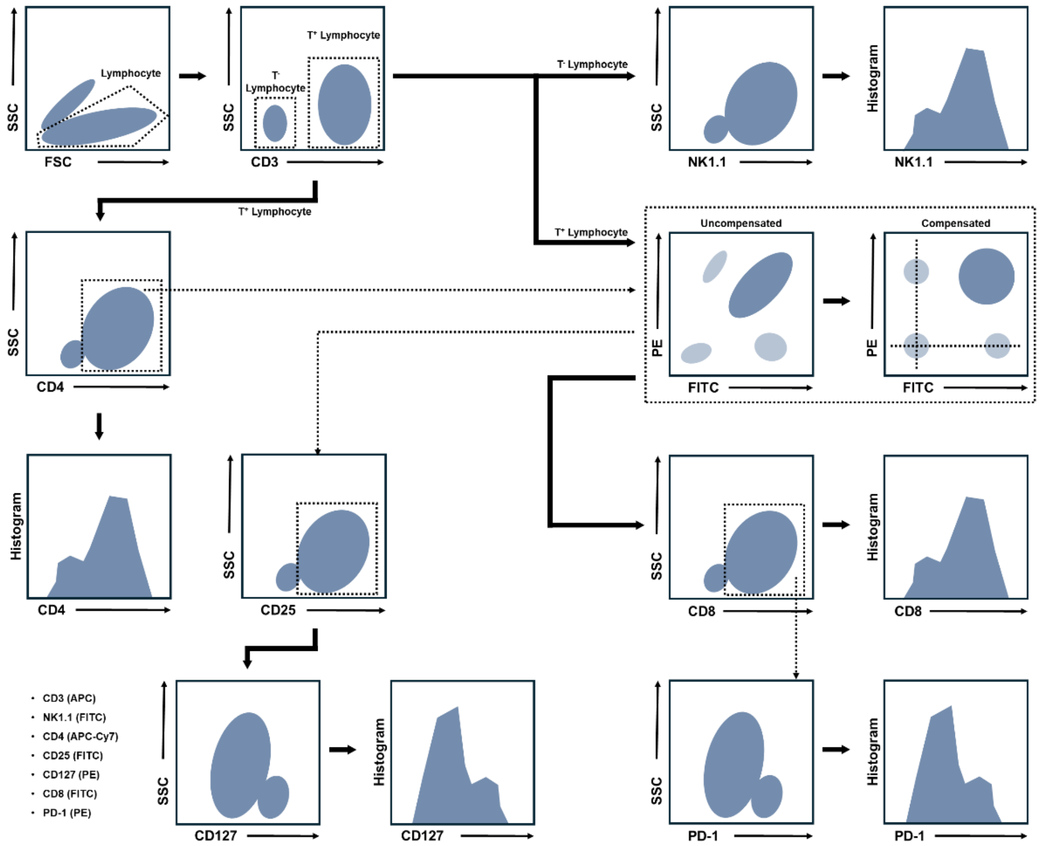


**Supplementary Fig. 41:** Workflow of flow cytometry analysis for TIME reprogramming test after mDC-sEV treatment. Percentage of the population was gated and quantified based on fluorescence intensity >5 × $10^3$ for NK1.1, CD4 and CD8; fluorescence intensity <5 × $10^3$ for CD127; fluorescence intensity <2 × $10^3$ for PD-1.

### SUPPLEMENTARY TABLES

| pH | mDC-sEVs | | mDC-sEVs(*DG*)/GALA | |
|---|---|---|---|---|
| | mean | s.e | mean | s.e |
| 7 | -15.325 | 1.440775 | -8.3925 | 1.502406 |
| 6 | -15.225 | 1.613227 | -3.83 | 0.559106 |
| 5 | -14.45 | 1.121011 | -0.00085 | 0.348145 |
| 4 | -14.9667 | 1.656301 | 1.197 | 0.334824 |

Unit: mV

**Supplementary Table 1:** Zetasizer analysis of GALA-functionalized mDC-sEVs under varying pH conditions (n = 4, mean ± s.e.).

96-well

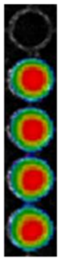

| Total Signal Intensity (Integrated Density; a.u.) | mean | s.d. |
|---|---|---|
| Ctrl (PBS) | $3.69\ e^{-5}$ | $6.39\ e^{-5}$ |
| 3T3-sEV | 1 | 0.004777 |
| imDC-sEV | 0.990528 | 0.016606 |
| mDC-sEV | 1.034803 | 0.0199 |
| mDC-sEV/GALA | 1.02072 | 0.00852 |

**Supplementary Table 2:** Fluorescence intensity of DiD-labeled sEVs measured in a 96-well plate and normalized to 3T3-sEV, used for quantification of *ex vivo* biodistribution (excitation/emission = 605/670 nm).

### AUTHOR INFORMATION


#### Corresponding Author

Yichun Wang - Department of Chemical and Biomolecular Engineering, University of Notre Dame, Notre Dame, Indiana 46556, United States; orcid.org/0000-0002-4353-6660; Tel: 574-631-2617;

Xin Lu - Department of Biological Sciences, University of Notre Dame, Notre Dame, Indiana 46556, United States; Tel: 574-631-6592;

#### Authors

Gaeun Kim - Department of Chemical and Biomolecular Engineering, University of Notre Dame, Notre Dame, Indiana 46556, United States; orcid.org/0009-0008-1502-1847

Shiyu Wang - Department of Biological Sciences, University of Notre Dame, Notre Dame, Indiana 46556, United States

Runyao Zhu - Department of Chemical and Biomolecular Engineering, University of Notre Dame, Notre Dame, Indiana 46556, United States; orcid.org/0009-0005-5109-0020

Matthew J. Webber - Department of Chemical and Biomolecular Engineering, University of Notre Dame, Notre Dame, Indiana 46556, United States; orcid.org/0000-0003-3111-6228

LYMPHOCYTES-IN-GLIOBLASTOMA-ARE.
